# A Constrained Mixture Theory Model to Study Autoregulation in the Coronary Circulation

**DOI:** 10.1101/2020.09.21.304030

**Authors:** Hamidreza Gharahi, Daniel A. Beard, C. Alberto Figueroa, Seungik Baek

## Abstract

Coronary autoregulation is a short-term response manifested by a relatively constant flow over a wide range of perfusion pressures for a given metabolic state. This phenomenon is thought to be facilitated through a combination of mechanisms, including myogenic, shear dependent, and metabolic controls. The study of coronary autoregulation is challenging due to the coupled nature of the mechanisms and their differential effects through the coronary tree. In this paper, we developed a novel framework to study coronary autoregulation based on the constrained mixture theory. This structurally-motivated autoregulation model required calibration of anatomical and structural parameters of coronary trees via a homeostatic optimization approach using extensive literature data. Autoregulation was then simulated for two different coronary trees: subepicardial and subendocardial. The structurally calibrated model reproduced available baseline hemodynamics and autoregulation data for each coronary tree. The autoregulation analysis showed that the diameter of the intermediate and small arterioles varies the most in response to changes in perfusion pressure. Finally, we demonstrated the utility of the model in two application examples: 1) response to drops in epicardial pressure, and 2) response to drug infusion in the coronary arteries. The proposed structurally-motivated model could be extended to study long-term growth and remodeling in the coronary circulation in response to hypertension, atherosclerosis, etc.

**Key points:**

- Coronary autoregulation is defined as the capability of the coronary circulation to maintain the blood supply to the heart over a range of perfusion pressures. This phenomenon is facilitated through intrinsic mechanisms that control the vascular resistance by regulating the mechanical function of smooth muscle cells. Understanding the mechanisms involved in coronary autoregulation is one of the most fundamental questions in coronary physiology.
- This paper presents a structurally-motivated coronary autoregulation model that uses a nonlinear continuum mechanics approach to account for the morphometry and vessel wall composition in two coronary trees in the subepicardial and subendocardial layers.
- The model is calibrated against diverse experimental data from literature and is used to study heterogeneous autoregulatory response in the coronary trees. This model drastically differs from previous models, which relied on lumped parameter model formulations, and is suited to the study of long-term pathophysiological growth and remodeling phenomena in coronary vessels.

## 1. Introduction

Coronary blood flow rapidly changes to meet changes in metabolic demand (myocardial oxygen consumption, MVO2) (25). This coronary flow regulation is the result of vessel diameter modulation via local control of the active stress in the smooth muscle cells (SMCs) (31). One manifestation of coronary flow regulation is the so-called pressure-flow autoregulation, in which perturbations in coronary perfusion pressure are met with commensurate changes in microvascular resistance to maintain a constant coronary blood flow at a given level of MVO2 (20).

Our current understanding of the different control mechanisms involved in coronary autoregulation is rather phenomenological. Briefly, coronary autoregulation relies on SMCs reactivity regulated through myogenic, shear-dependent, and metabolic control mechanisms. Myogenic control is the intrinsic response of SMCs to changes in the local wall stress (52). Shear-dependent control is a dilatory mechanism mediated via shear-induced production of nitric oxide (NO) by the endothelial cells. Metabolic control is a local feedback mechanism where a vasodilation signal is initiated in response to either an increase in local oxygen consumption or oxygen extraction from the myocardial blood supply (25). Studies have shown that these mechanisms heterogeneously affect the coronary microvasculature (81). Myogenic control has been observed to be small in precapillary arterioles and large in the arteries of 50-150 μm (55). Shear-dependent control is most active in the arteries and larger arterioles (54). Lastly, the signal for the metabolic feedback mechanism is initiated in the capillaries, conducted upstream, and it gradually decays for larger vessels (∼150 μm) (44). Furthermore, coronary autoregulation occurs across different layers of the myocardium where the myocardial-vessel interactions are not uniform (92). Therefore, the anatomical and functional heterogeneity of the coronary arteries, and the difficulties in acquiring *in-vivo* measurements necessitate the development of computational models that can integrate the available experimental data on morphometry, biomechanical, and physiological responses to enhance our understanding of coronary autoregulation in healthy, disease, and treatment planning conditions (e.g., coronary artery disease, cardiac hypertrophy, metabolic dysfunction, etc.) (23).

Several computational techniques have been used to model the morphometry and hemodynamics of coronary arteries (1, 12, 39, 46, 77). Kaimovitz et al. (42, 43) provided a stochastic framework for constructing the entire coronary arterial vasculature using a volume-filling optimization subject to morphometric statistics reported in (48). Namani et al. (67) extended Kaimovitz’s model by incorporating physiological hemodynamic constraints in the generation of the coronary arterial network. They showed that a concurrent minimization of flow dispersion and diameter re-assignment at the microvascular level resulted in a realistic representation of the network when compared with literature data. These coronary arterial tree reconstruction methods, while informed by morphometry and hemodynamic data, have not considered structurally-motivated data for the passive and active behavior of the coronary vessels at the different generations of the network.

On the other hand, the phenomenon of coronary autoregulation has been the subject of numerous modeling studies over the last 50-60 years. Virtually all of such studies have relied on lumped-parameter (0D) approaches. Liao and Kuo (55) and Cornelissen et al. (18) investigated the interaction and balance between autoregulatory mechanisms in the coronary circulation. While these studies incorporated the heterogeneity of the microvascular response, they considered a simple partitioning of the coronary tree into 4 compartments and ignored the interactions between vessels and myocardium. Pradhan et al. (69) applied a data-driven closed loop model of coronary flow regulation using *in-vivo* data on coronary flow and oxygen extraction in response to exercise-induced changes in demand and to changes in perfusion pressure. The Pradhan model, which represents parallel control pathways using a block-diagram approach, does not represent explicit spatial features. More recently, Namani and colleagues (66) developed a model of the coronary microcirculation that integrated the dynamic effects of coronary flow with myogenic, shear dependent, and metabolic feedback control mechanisms in subendocardial and subepicardial vessels. Their constitutive model considered simple pressure-diameter rules to define the behavior of the vessels. While the aforementioned studies provide reasonable descriptions of autoregulatory responses, they lacked structurally-motivated models for arterial tissue. Given that the short-term regulation of vascular tone and the long-term vascular growth and remodeling processing, including pathophysiologic responses, are highly intertwined (85), there is a pressing need to develop models of coronary autoregulation that can account for the microstructure of the vessel wall.

Constrained mixture theory models have been widely applied to describe the nonlinear mechanical behavior of arterial tissue, accounting for the contributions of the main load-bearing constituents (e.g., collagen, elastin and SMCs) (26, 74). This theory provides a formal means to represent mechanical function of a vessel based on the properties of its constituents and the relative mass fractions of those constituents in the composition of the vessel wall. Using this theory, long-term growth and remodeling processes are represented by changes in the composition of the vessel wall. Previous application of the constrained mixture theory in vascular mechanics have focused on large arteries where homeostatic stress has been assumed constant. However, wall stress has been found to be size-dependent through the vasculature (33, 71). Motivated by this, Filonova and colleagues (29) developed a homeostatic optimization framework, adopted from an extended Murray law (56, 80), to generate vascular trees obeying a data-based fractal, to predict homeostatic characteristics, such as morphometry, hemodynamics, and structure.

The goal of this study is to apply the homeostatic optimization framework of Filonova et al. to develop a structurally-motivated computational model of coronary autoregulation, a physiological mechanism that fundamentally relies on mechanical function of SMCs. The model, based on the constrained mixture theory, provides a means to estimates baseline activation of SMCs and homeostatic stresses, while being informed by experimental pressure-diameter, tree morphometry, and hemodynamics data, for both subepicardial and subendocardial trees. In this work, our model is limited to calculating the steady-state response to stimuli triggering combined alterations in metabolic, shear-dependent, and myogenic mechanisms. Lastly, the utility of our model is demonstrated using two application examples: 1) study of short-term adaptations following drops in epicardial pressure (Section 3.4), and 2) simulation of the short-term coronary responses to pharmacological agents such as adenosine (which binds to purinergic receptors and causes SMC relaxation) and L-NAME infusion (which inhibits NO synthesis and therefore induces vasoconstriction) (Section 3.5). An additional benefit of our approach is that this model could also be extended to study long-term growth and remodeling in the coronary circulation in response to hypertension, atherosclerosis, etc., thereby rendering a unified framework to describe the dynamics of both short-and long-term coronary vascular adaptations.

## 2. Methods

### 2.1. Model overview and parameter estimation process

Coronary arterial networks embedded in the subendo-and subepicardial layers are represented as bifurcating trees. A steady state fluid model describes the relation between blood flow and pressure through estimation of vascular resistance in each idealized tree segment. The geometry and composition of vessel segments at each generation of the tree are defined based on minimization of energy consumption and the mechanical equilibrium which relate the diameter, pressure, and vessel wall thickness. The model uses a nonlinear continuum mechanics approach and is endowed with autoregulatory mechanisms to study the coronary autoregulation. Overall, our model determines the diameter of each vessel segment as a function of the coronary perfusion pressure by integrating the hemodynamics of the tree and autoregulatory responses.

The strategy to calibrate the model is summarized in **Fig. 1**. The first stage is to estimate the constrained mixture model parameters, representing the material properties of load-bearing constituents within the vessel wall (elastin, collagen, SMCs), their respective pre-stretches (the stretch at which the constituents are deposited in homeostatic conditions), and their mass fractions, for four vessel types (small arteries, large, intermediate, and small arterioles), for both the subepicardium and subendocardium layers. These parameters are estimated by comparing model predictions to data on measurements of the passive and full myogenic response of isolated porcine coronary vessels to internal pressurization (55). In addition to the constrained mixture model parameters, this step also identifies model parameters associated with the myogenic component of the autoregulatory response.

**Figure 1.**
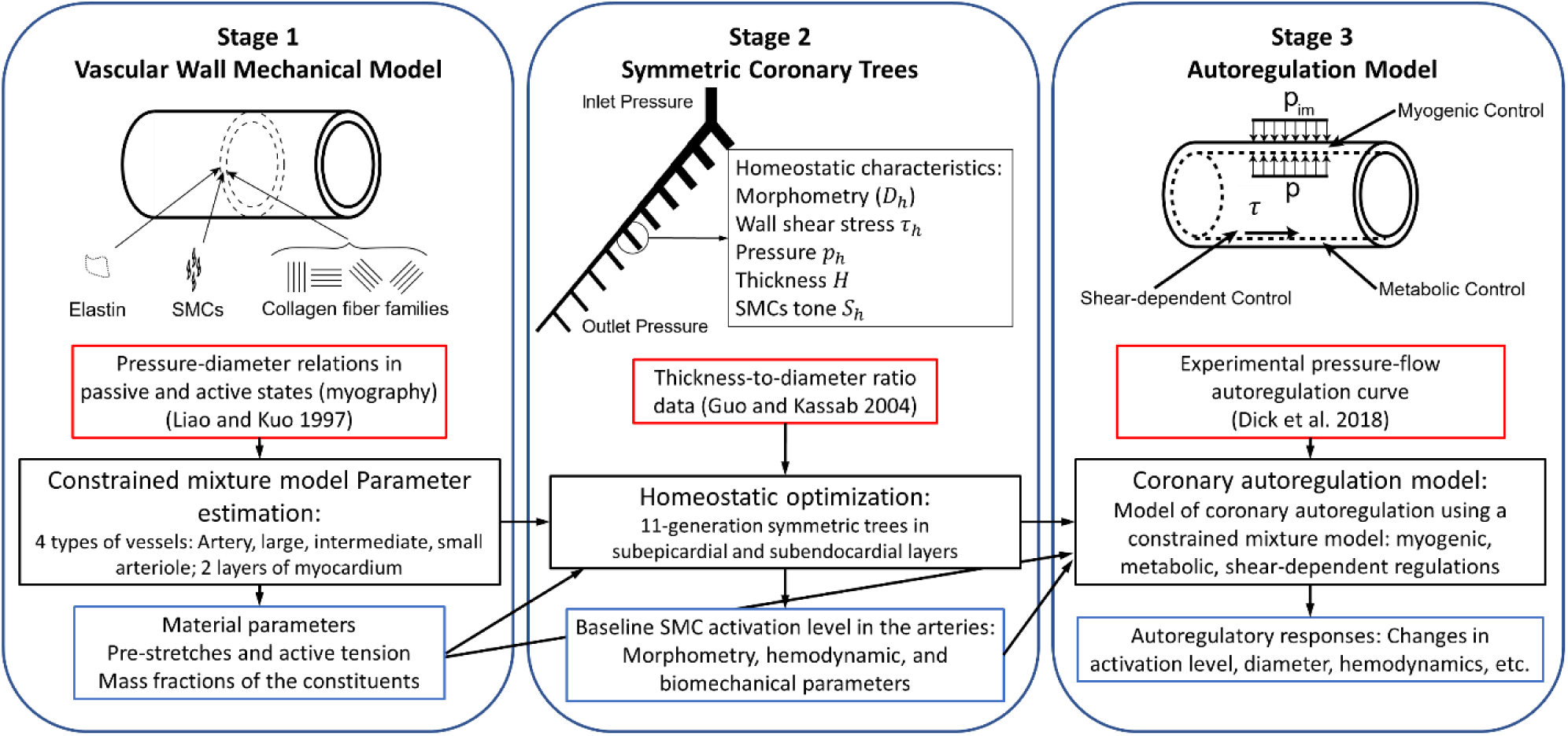
Schematic of the components and the workflow of modeling stages and data in the proposed constrained mixture theory model of coronary autoregulation. The arrows show the flow of information. Red, black, and blue boxes delineate the literature data, the methods used, and model results, respectively.

Second, using the homeostatic optimization framework proposed by Filonova and colleagues (29), we generate two coronary trees in the subendocardial and subepicardial layers and establish their homeostatic characteristics, i.e., morphometry, hemodynamics, and structure. Motivated by Murray’s law (63), the homeostatic optimization framework defines the homeostatic state (equilibrium) of a vascular tree to be governed by an energy-based minimization problem in which the energy dissipation includes the power needed to overcome viscous drag and the metabolic energy spent on sustaining blood supply and maintaining the constituents of the vascular wall. This minimization is performed under mechanical equilibrium as a constraint.

Third, a structurally-motivated coronary autoregulation model was developed, integrating the myogenic with metabolic and shear-dependent mechanisms, for each of the two coronary trees. In the following sections, we describe each of the three stages in detail, together with the experimental data they require.

### 2.2. Parameter estimation for the constrained mixture model of 8 representative coronary vessels

A constrained mixture theory model is used to describe the arterial tissue mechanics with three main load-bearing constituents: elastin matrix, collagen fibers, and SMCs. This approach incorporates microstructural properties and cellular level functions of the vessel within a nonlinear continuum mechanics approach, whereby the wall constituents are constrained to deform together but have mechanical properties and stresses. The relationship between transmural pressure *p*_*tm*_ and internal diameter *D* in a thin walled cylinder is given by Laplace’s law: *p*_*tm*_*D*/2 = *T*_*θθ*_, where the circumferential tension *T*_*θθ*_ is determined by contributions from elastin, collagen and SMCs. Each constituent has distinct strain energy densities (*w*) mapping deformation gradients from their distinct stress-free configurations to the overall *in-vivo* configuration. In this work, we considered a neo-Hookean model for elastin (with coefficient *c*_1_) and a Holzapfel’s exponential model for i) collagen fiber families (with coefficients *c*_2_ and *c*_3_) and ii) passive response of SMCs (*c*_4_ and *c*_5_). The active tension (*T*_*act*_) exerted by SMCs, was defined as

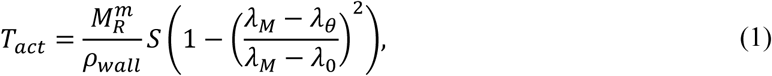

where *S* is the active SMC stress, *ρ*_*wall*_ is the density of arterial tissue, 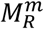 areal density of SMCs, *λ*_*M*_ and *λ*_0_ are stretches corresponding to the maximum and zero active tension, respectively, and *λ*_*θ*_ is the vessel circumferential stretch. Since details of general mathematical forms of passive and active behaviors of the constrained mixture theory models are available elsewhere (10), they are briefly summarized in **Appendix A**.

The three main control mechanisms controlling active SMC stress *S* in coronary autoregulation are myogenic, shear-dependent, and metabolic. In this work, we are interested in a formulation that separates the myogenic contribution from the shear-dependent and metabolic contributions to SMC stress *S*. Therefore, we have assumed the following functional relationship for *S:*

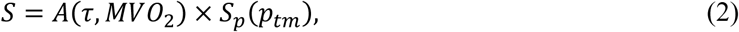

where *A* is the activation level (0 < *A* < 1), determined by wall shear stress (*τ*) and metabolic activity (*MVO*_2_) mechanisms; and *S*_*p*_ is a pressure-dependent stress representing the level of myogenic control. Thus, when *A* = 0, a given vessel is in a maximally dilated state, regardless of any influence of the myogenic mechanism. When *A* = 1, there is no vasodilation, and the control of smooth muscle activation is exclusively through the myogenic mechanism. Following Cornelissen and colleagues (18), we assume that *S*_*p*_ follows a Hill curve for positive transmural pressures and is zero in negative transmural pressures:

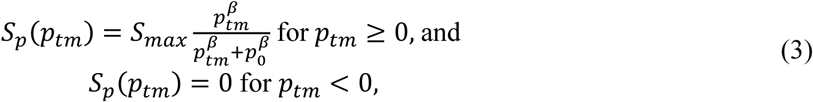

where *β* is the Hill slope, *S*_*max*_ is the total maximum active stress, and *p*_0_ is a parameter that offsets the center of the curve.

In addition to the material parameters determining the active and passive behavior of individual constituents, other important parameters that define the mechanical homeostatic condition in the constrained mixture models of vascular tissue are: 1) the homeostatic pre-stretches at which each constituent is deposited (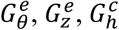, and 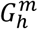, see **Appendix A** for details), and 2) the fraction of the vessel wall occupied by each constituent (mass fractions *v*^*e*^, *v*^*c*^, and *v*^*m*^; *v*^*e*^ + *v*^*c*^ + *v*^*m*^ = 1).

Constrained mixture models have been widely used in the computational study of growth and remodeling of large arteries, under the general assumption that the homeostatic stress of each wall constituent is constant (2, 28, 30, 91). However, the homeostatic stress of each constituent is unlikely to remain constant along the tree, since stresses such as hoop stress and wall shear stress vary significantly from large arteries to small arterioles (29, 71). Therefore, to account for this likely variability in constituent homeostatic stress, in this work we assumed that the mechanical properties of constituents (material stiffnesses *c*_1_−*c*_5_ and active SMC parameters *S*_*max*_, *β, p*_0_, *λ*_*M*_, *λ*_0_) are constant across the vascular tree, but that the parameters representing the wall composition (mass fractions) and homeostatic deposition stretch of the constituents (pre-stretches) are vessel-specific (i.e., vary with vessel size and its homeostatic transmural pressure). **Table 1** summarizes the parameters of the constrained mixture model and their definitions.

**Table 1.** Parameters defining the mechanical properties of the vessel wall model. The passive and active material parameters are constant across all representative vessels. The vessel-specific parameters vary between representative vessels with respect to their size and myocardial location.

|  | Parameter | Description | Unit |
| --- | --- | --- | --- |
| Passive material parameters<br>(Appendix A) | $c_1$ | Elastin material parameter | Pa/kg |
| | $c_2$ | Collagen material parameter | Pa/kg |
| | $c_3$ | Collagen dimensionless material parameter | - |
| | $c_4$ | SMCs material parameter | Pa/kg |
| | $c_5$ | SMCs dimensionless material parameter | - |
| Active SMC model parameters<br>(Section 2.1) | $S_{max}$ | Total maximum stress | MPa |
| | $\beta$ | Hill curve slope | - |
| | $p_0$ | Hill curve offset | mmHg |
| | $\lambda_0$ | Zero active tension stretch | - |
| | $\lambda_M$ | Maximum active tension stretch | - |
| Vessel-specific parameters<br>(Appendix A) | $G_\theta^e, G_z^e$ | Elastin pre-stretch | - |
| | $G_h^c$ | Collagen pre-stretch | - |
| | $G_h^m$ | SMCs pre-stretch | - |
| | $v^e$ | Elastin mass fraction | - |
| | $v^c$ | Collagen mass fraction | - |
| | $v^m$ | SMC mass fraction | - |

#### Parameter estimation

Liao and Kuo (55) measured the passive and myogenic pressure-diameter relations in four classes of swine coronary vessels with internal diameters of 180 ± 20 μm (small arteries), 95 ± 2 μm (large arterioles), 62 ± 2 μm (intermediate arterioles), and 37 ± 2 μm (small arterioles) using pressure myography. In this work, we used these relations to estimate the parameters of the constrained mixture model for the four representative vessel types in each subepicardial and subendocardial layers. To replicate the passive pressure-diameter relations, we fixed the shear-dependent and metabolic activation level and myogenic control to zero (*A* = 0 and *S*_*p*_ = 0). Conversely, to mimic the myogenic data of Liao and Kuo, we assumed no contribution to vasodilation from either the shear-dependent or the metabolic mechanisms, i.e., *A* = 1. In this case, the active SMC stress *S* is given entirely by the myogenic control *S*_*p*_ (Eq. 3).

Overall, identifying the parameters of the constrained mixture model of the eight representative vessels required estimating a total of 50 parameters. Several physiological assumptions and constraints were considered to avoid unrealistic values of the model parameters. The collagen fiber angles were prescribed to follow the axial, circumferential, and diagonal directions (e.g. 0°, 90° and 45°, respectively). Collagen is assumed to have a relatively low pre-stretch, thus we used 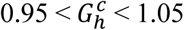. The pre-stretches of elastin 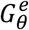 and 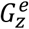 and SMCs 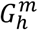, however, were assumed to be greater than 1 (84).

The homeostatic condition of a vessel, i.e., its *in-vivo* diameter, thickness, and transmural pressure, is needed to identify the parameters of the constrained mixture model. In this work, we assumed an *in-vivo* vessel diameter (*D*_*h*_) corresponding to the experimental data (55), and obtained the vessel-specific pressure (*p*_*h*_) and thickness (*H*) from literature (33, 62). Finally, the distension pressure in the Liao and Kuo experiments was always positive, and thus, the data did not describe the vessel response under compression. Therefore, in this work we assumed that the active SMC stress *S* under compression is zero, similar to findings in skeletal muscle circulation (17). The downhill simplex method was used to estimate the parameters of the constrained mixture model (76) by minimizing the mean squared error (*E*) between predicted and measured diameters for a given transmural pressure.

The *in-vivo* activation level *A* for a given vessel should be in the range of 0 and 1. Therefore, once the parameters associated with passive (*A* = 0 and *S*_*p*_ = 0), and myogenic (*A* = 1) responses are identified, an *in-vivo* vessel-specific homeostatic (baseline) activation *A*_*h*_ can be computed using *D*_*h*_ and *p*_*h*_. Consequently, the homeostatic active SMC stress *S*_*h*_can be calculated via Eq. 2: *S*_*h*_ = *A*_*h*_ × *S*_*p*_(*p*_*h*_).

Once the parameters of the constrained mixture model (**Table 1**) for the representative vessels (4 in subendocardial and 4 in subepicardial layer) are identified, they are assigned to each individual vessel of the coronary trees according to its size.

#### Sensitivity analysis

To understand the sensitivity of the estimated parameters of the constrained mixture model, a sensitivity analysis was performed following the approach of Pradhan and colleagues (69). A sensitivity index *X* was defined as the percentage change in model error *E* (**Appendix B**) associated with introducing a 10% change in the parameter values one at a time

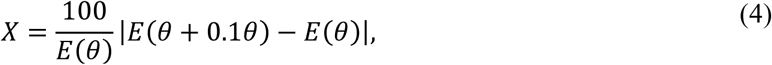

where *θ* is the estimated parameter value, and *E*(*θ*) is its associated error, defined as the difference between predicted and measured diameters.

### 2.3. Defining morphometry, structure, and hemodynamics of an entire coronary tree

The second stage is to generate two coronary trees in the subendocardial and subepicardial layers and to estimate their homeostatic baseline state, using the homeostatic optimization framework developed by (29). The approach of Filonova et al. is a framework to estimate the homeostatic morphometric (e.g, diameters), structural (e.g, thickness), and hemodynamics (e.g., wall shear stress) properties of a vascular tree by minimizing the global energy dissipation under the constraint of mechanical equilibrium. This energy minimization is performed via adjustments of 2 parameters in each vessel segment of the tree: 1) the homeostatic diameter of the vessel and, 2) the amount of the materials of the vessel wall that constitute the thickness (elastin, collagen, SMCs) (**Appendix C**).

#### Tree initialization

First, two symmetric trees (subendocardial and subepicardial) were initialized using morphometry data. The parent-to-daughter relationship is used to describe the branching pattern of the vascular trees

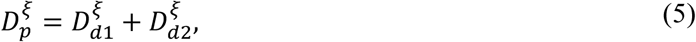

where *D*_*p*_, *D*_*d*1_, and *D*_*d*2_ are the diameters of the parent and two daughter vessels, respectively, and *ξ* is called the diameter exponent. Vessels feeding the intramyocardial arteries of the left ventricular free wall are 438±56 μm in diameter (42). With an initial radial exponent of *ξ*_0_ = 2.55 (46), the diameter of the first-generation vessel equal to 438 μm and the diameter of the last generation vessels ∼ 20 μm, the number of generations in the trees is 11. Vessel lengths were determined as a function of vessel diameters using the morphometric data in Kassab et al. (48). The subendocardial and subepicardial trees were assumed to be located at 5/6 and 1/6 of the myocardial depth, respectively.

#### Tree optimization

We considered hemodynamic steady state conditions to define baseline homeostatic characteristics of the trees, with distinct mean intramyocardial pressures for each tree. Using a thin-walled assumption, the mechanical equilibrium in each artery can be expressed via Laplace’s law:

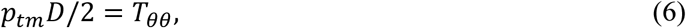

where *p*_*tm*_ is the transmural pressure of the myocardial blood vessels, *T*_*θθ*_ is the arterial membrane stress that incorporates the mechanical contributions of elastin, collagen, and SMCs (**Appendix A**), and *D* is the inner diameter of the segment. The transmural pressure can be written as

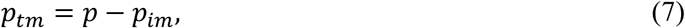

where *p* is the luminal pressure and *p*_*im*_ is the intramyocardial pressure. Algranati and colleagues (3) postulated that *p*_*im*_ is a function of the cavity-induced extracellular pressure (*p*_*CEP*_) and muscle shortening-induced pressure (*p*_*SIP*_):

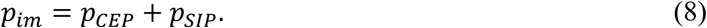

Dependence on *p*_*CEP*_ implies that the compressive forces are larger in the inner layers (endocardial) of the left ventricle (LV) than those in the outer (epicardial) layers (7). Furthermore, it has been established that this variation in myocardial pressure affects the flow distribution and structure of the arteries in different layers (50). We assumed that for the free wall of the LV, *p*_*CEP*_ varies linearly from endocardium (LV pressure, *p*_*LV*_) to epicardium, where the pericardial pressure is assumed to be negligible. Therefore, for the subendocardial and subepicardial trees, the average *p*_*CEP*_ values were set to 5/6 and 1/6 of *p*_*LV*_, respectively, consistent with their relative myocardial depth. The muscle shortening-induced pressure *p*_*SIP*_ was chosen so that subendocardium *p*_*im*_ is ∼20% larger than the *p*_*LV*_ (65).

The boundary conditions (i.e., hemodynamic constraints) of the homeostatic optimization in both subepicardial and subendocardial trees are the inlet and outlet pressures. In general, imposing only pressure boundary conditions is not sufficient for the homeostatic optimization as the solution will not converge (29). Therefore, a myocardial layer-dependent total flow in the arterial tree was considered as a soft constraint for the global optimization, by adding a penalty term to the cost function (**Appendix C**). A downhill simplex method was used for the minimization of the global cost function. The direct result of the minimization is the diameter of the vessels in coronary trees. Once the vessel diameter is known, the vessel wall thickness can be computed as a post-processing using the mechanical equilibrium equation (**Appendix A**).

#### Model parameters

In this work, the bulk of model parameters corresponds to literature swine data, with the exception of flow at terminal arterioles, which corresponds to canine data, given lack of measurements for swine (67). Next, we detail the data used to set our hemodynamics and how the constrained mixture model parameters are assigned to the coronary tree vessels.

The hemodynamics parameters used for generating the coronary trees are listed in **Table 2**. Baseline coronary pressure was taken to be the mean aortic pressure 100 mmHg since the pressure drop in epicardial arteries, in absence of occlusive disease, is negligible. The outlet pressure at the terminal vessels (small arterioles) was set to 55 mmHg (27, 72). Based on blood flow velocity measurements in (8, 78, 82), a baseline flow rate of 3 × 10^−3^ mm^3^/s was used for the subepicardial terminal arterioles. To determine the flow for the subendocardial tree, the total subendocardial to subepicardial flow ratio (ENDO/EPI) was assumed to be ∼1.25 (22). Intramyocardial pressures *p*_*im*_ of 47 and 13 mmHg were imposed on subendocardial and subepicardial vessels, respectively. A diameter dependent blood flow viscosity model was used based on (73) (**Appendix D**).

**Table 2.** Tree initialization and hemodynamic parameters for generation of coronary trees.

| Parameter | Description | Value | Unit | Ref |
| --- | --- | --- | --- | --- |
| $\xi_0$ | Initial diameter exponent | 2.55 | - | (46) |
| $D_0$ | Initial first-generation diameter | 438 | $\mu\text{m}$ | (42) |
| $p_{in}$ | Inlet coronary pressure | 100 | mmHg | |
| $p_{out}$ | Terminal pressure | 55 | mmHg | (27, 72) |
| $q_{subepi}$ | Subepicardial (baseline) terminal flow | $3 \times 10^{-3}$ | $\text{mm}^3/\text{s}$ | (8, 78, 82) |
| $ENDO/EPI$ | Subendocardial to subepicardial flow ratio | 1.25 | - | Duncker and Bache 2008) |
| $p_{im}$ (subepi., subendo.) | Intramyocardial pressure | 13, 47 | mmHg | (65) |

Following (18), the constrained mixture model parameters were assigned to the coronary vessels according to their sizes (Arteries: D>190μm, Large Arterioles: 190≥D>100 μm, Intermediate Arterioles: 100≥D>50, Small Arterioles: 50≥D) for each tree. This classification is limited by the experimental data and is relatively crude since the pressures and wall thicknesses within one class of vessels may vary significantly. For instance, our initial analysis showed that when all the parameters are constant across a class type, the homeostatic optimization framework does not produce the correct thickness-to-diameter ratio trends when compared to experimental data. Therefore, we adjusted the baseline activation level of SMCs (0 < *A*_*h*_ < 1), which is the *in-vivo* value of *A* in Eq. 2, for each vessel to match the thickness-to-diameter ratio with data in (33), as demonstrated in the results section.

### 2.4. Coronary autoregulation model

The third stage of the proposed model is to develop a structurally-motivated coronary autoregulation model based on constrained mixture theory. In coronary autoregulation, vessel reactivity to changes in pressure is mediated by the active SMC stress *S* (Eq. 2). Although autoregulation is a dynamic process involving the activation of each of the metabolic, shear-dependent and myogenic mechanisms with their respective time responses, in this work we are concerned with the steady state reached within minutes following a perturbation from a previous state (5, 19, 66).

A key component of the proposed autoregulation model is the activation function *A* (Eq. 2), which determines the activation level from a fully dilated (*A* = 0) to a maximum myogenic response (*A* = 1). The myogenic control *S*_*p*_, is a constrictive response to increases in the local wall stress resulting from changes in transmural pressure *p*_*tm*_. The myogenic control is formulated by the sigmoidal function in Eq. 3.

In shear-dependent control, an increase in wall shear stress *τ* induces relaxation of the SMCs mediated by upregulation of endothelial NO synthesis. This vasodilatory stimulus 𝓈_*τ*_ is modeled as

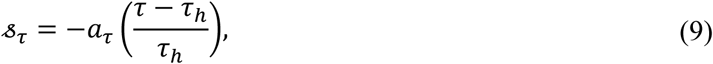

where *τ*_*h*_ is the baseline (homeostatic) value of wall shear stress, and *a*_*τ*_ is the shear-dependent scaling coefficient. The negative sign indicates a decrease in vessel activation for an increase in *τ*.

Moreover, experimental studies show that the oxygen extraction capacity of the myocardium is almost fully exhausted in both rest and exercise (24). Therefore, we assume that changes in *MVO*_2_ are matched by proportional changes in flow to supply the oxygen demand (6). Based on this assumption, we take the terminal flow as a measure of metabolic signal, regardless of the underlying physiological mechanisms involved (66), and write the metabolic stimuli as

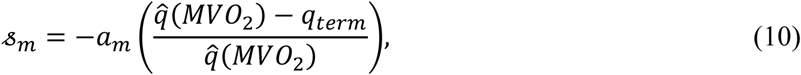

where 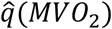 is the target flow determined by baseline metabolic demand, *q*_*term*_ is the actual flow rate at the terminal arterioles, and *a*_*m*_ is the metabolic scaling coefficient. The negative sign indicates a decrease in metabolic stimuli for a decrease in terminal flow. It must be noted that in this work, we focus on the coronary autoregulation for a fixed baseline myocardial activity, i.e., 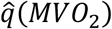 remains constant.

To compute the total stimuli induced by the shear-dependent and metabolic control mechanisms, two factors must be considered. First, the magnitude of each control mechanism depends of the vessel size. Shear-dependent vasodilation is mostly blunted in vessels smaller than 100 μm (78). Meanwhile, the metabolic response from different signaling pathways (e.g., oxygen demand/supply) is initiated in capillaries, and it is conducted and integrated upstream to precapillary arterioles (5). The integrated response, however, decays exponentially so that it mostly affects the intermediate and small arterioles with a peak at vessels ∼50 μm. Second, the equations 9 and 10 only describe the stimuli when a deviation from homeostatic conditions occurs. However, SMCs maintain a basal tone under resting conditions (59). Therefore, the total SMC stimuli for each vessel can be expressed as a basal stimulus plus the superposition of the two stimuli described above, viz.:

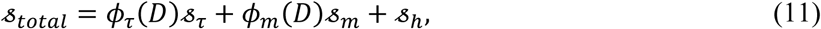

where 𝓈_*h*_ is a SMC basal (homeostatic) stimuli. Based on the experimental observations (13, 45, 53, 54), the diameter-dependent weights *ϕ*_*τ*_ and *ϕ*_*m*_, representing the magnitude of each mechanism in the total stimulation, are prescribed (**Fig. 2**)

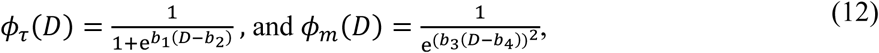

where *b*_1_ and *b*_3_ are 0.02 and 0.013 1/μm, and *b*_2_ and *b*_4_ are 250 and 50 μm, respectively. Following (15), a sigmoidal function is used to relate the total stimuli *s*_*total*_ to the activation function *A*:

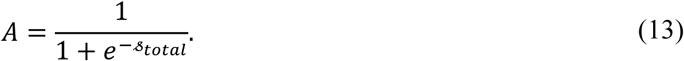

**Figure 2.**
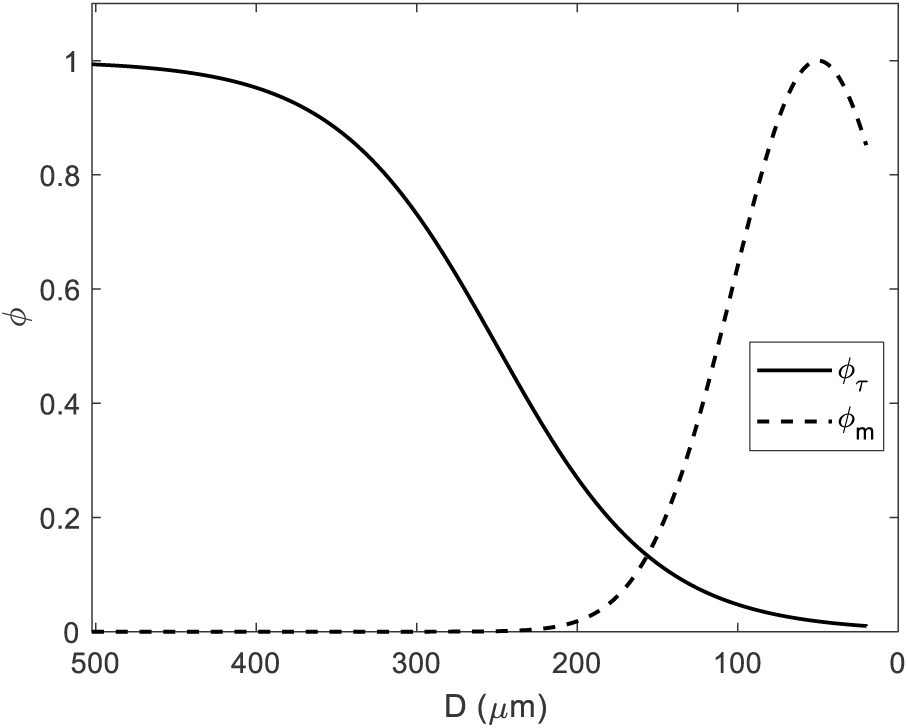
Magnitude of the shear-dependent and metabolic mechanisms, based on (13, 45, 53, 54). The shear-dependent mechanism is weak in arterioles smaller than 100 μm. A metabolic signal is initiated at the end of terminal vessels (capillaries/venules) and is conducted upstream to coronary arterioles, where signals from different terminal vessels are integrated. The metabolic signal, however, decreases exponentially for larger vessels. The magnitude of the metabolic mechanism was assumed to be highest at the arterioles (peak at 50 μm).

Therefore, the basal stimuli 𝓈_*h*_ for each vessel in the coronary tree can be calculated using the homeostatic activation level *A*_*h*_

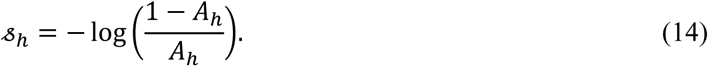

Combining equations (2)-(3),(9)-(13), we obtain expression for the total active SMC stress *S* as a function of *p*_*tm*_, *τ*, and *MVO*_2_.

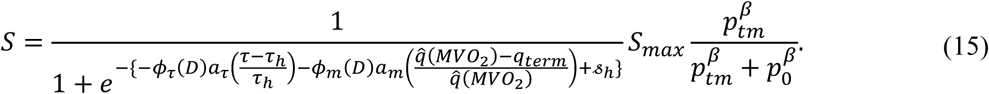

The scaling coefficients *a*_*τ*_ and *a*_*m*_ for the autoregulation model are estimated using the collection of swine coronary autoregulation data in (21). The procedure and the results of this calibration are described in Section 3.3. **Table 3** summarizes the model parameters for the autoregulation model and how the values are estimated/prescribed.

**Table 3.**
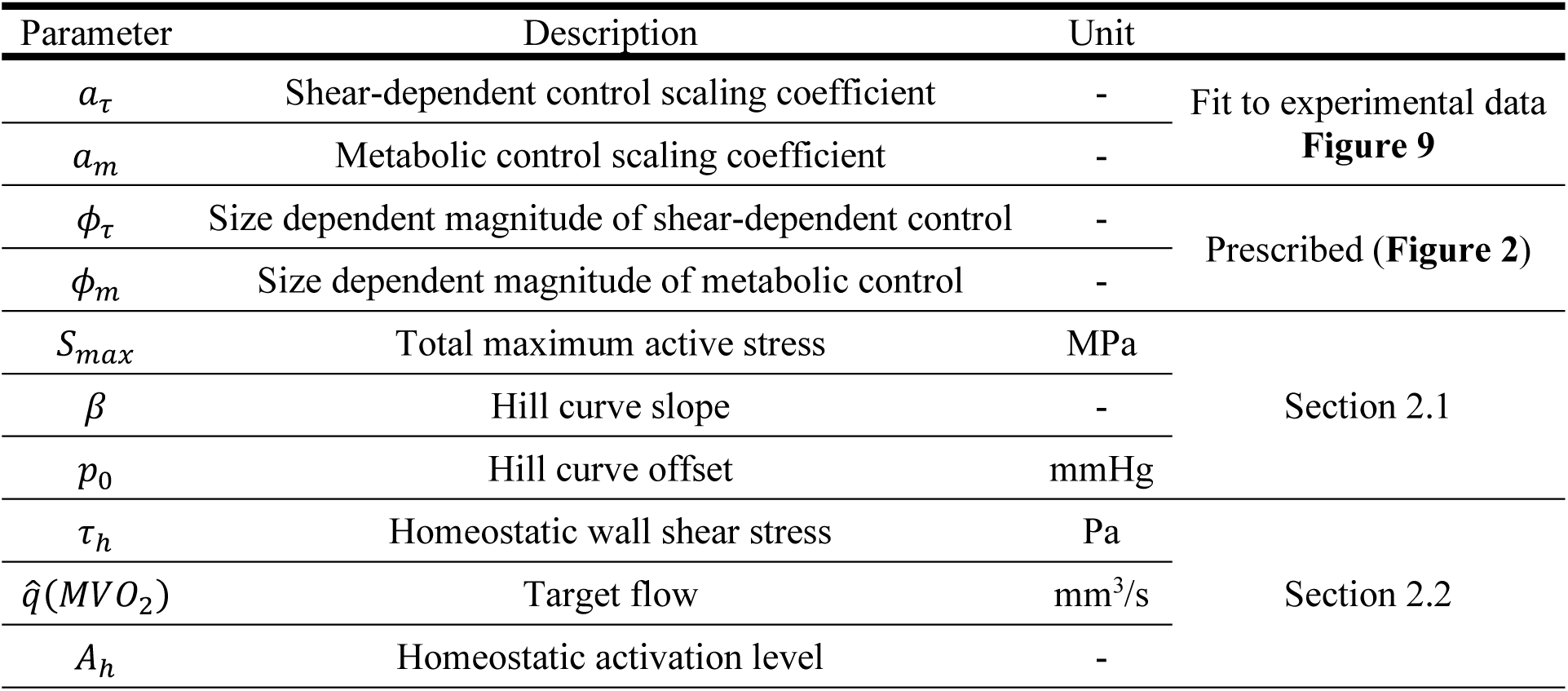
Autoregulation model parameters

The coronary pressure will therefore critically determine the relative contribution of the different control mechanisms down the coronary tree. Pressure and flow rates are calculated in each individual vessel of the entire coronary tree. A change in input coronary pressure elicits a new hemodynamic state and corresponding stimuli from each control mechanism. New SMC activation levels and active SMC stress are computed and used in the constrained mixture model to determine the changes in vessel size, which in turn result in an updated global hemodynamic state. This process is repeated until the tree hemodynamics are converged, which defines the new regulated state. To ensure the uniqueness of the converged steady state, the initial conditions for the global steady state hemodynamics were randomly varied. This workflow is depicted in **Fig. 3**.

**Figure 3.**
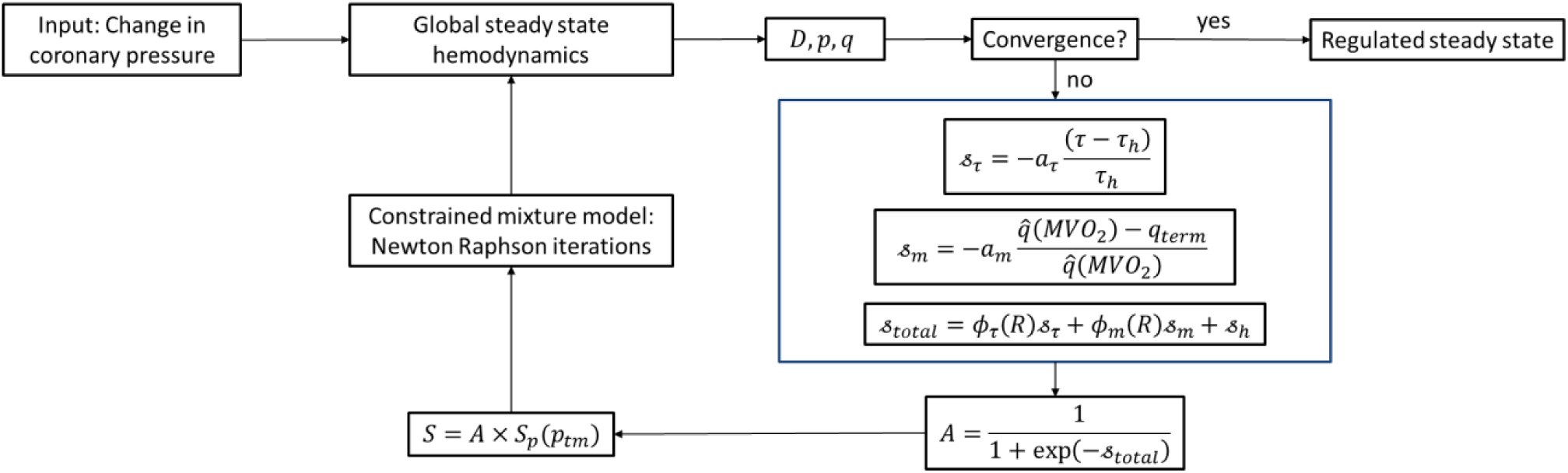
The workflow of the autoregulation model. The model input is the coronary pressure *p*_*in*_. Convergence in hemodynamics (pressure and flow in the entire tree) defines a new regulated state.

## 3. Results

In Sections 3.1 and 3.2, we present the results of model calibration for the constrained mixture model parameters and for the morphometry, structure, and hemodynamics of the different coronary trees, respectively. In Section 3.3, we illustrate the autoregulation model by calculating responses every 5 mmHg of input coronary pressure for the 20-160 mmHg pressure range. All the stages of the model were implemented in MATLAB (Mathwork, Natick, MA).

### 3.1. Constrained mixture model parameters

Table 4 and **Fig. 4** show the constrained mixture model parameter values estimated using the experiment data from Liao and Kuo (55) depicted in **Fig. 5**. Relatively higher mass fractions of SMCs were estimated in the large and intermediate arteriolar levels compared to small arteries and small arterioles for both coronary trees (**Fig. 4**). In addition, the pre-stretch of collagen fibers in the small arterioles of subepicardium and all arterioles of subendocardium were estimated to be less than 1, which means that these fibers are slightly compressed in the homeostatic condition of the vessel (**Table 4**). The SMC passive material parameters show the lowest sensitivity (see sensitivity indices in **Table 4**). In **Fig. 5**, the model results are compared with experimental data for passive (*A* = 0 and *S*_*p*_ = 0) and myogenic (*A* = 1 and *S*_*p*_ from Eq. 3) responses.

### 3.2. Morphometry, structure, and hemodynamics of an entire coronary tree

Morphometric results of the homeostatic optimization are summarized in **Fig. 6**. The homeostatic optimization estimates vessel diameters *D*_*h*_, determines bifurcation rules, identifies vessel types based on their sizes, and assigns the appropriate mechanical parameters (29). Subendocardial vessels appear to be larger than their subepicardial counterparts. This difference, consistent with experimental and modeling results from (3, 35), facilitates a higher blood flow in the subendocardial than in the subepicardial layer. Furthermore, our results indicate an increasing diameter exponent *ξ* (2.5-2.7) with generation number for both trees.

**Table 4.**
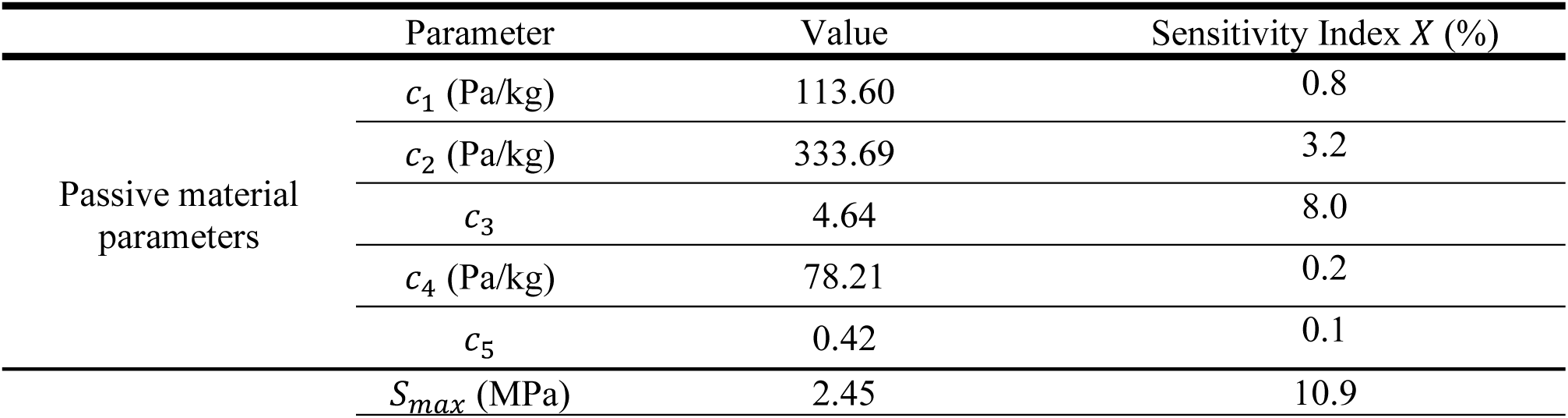
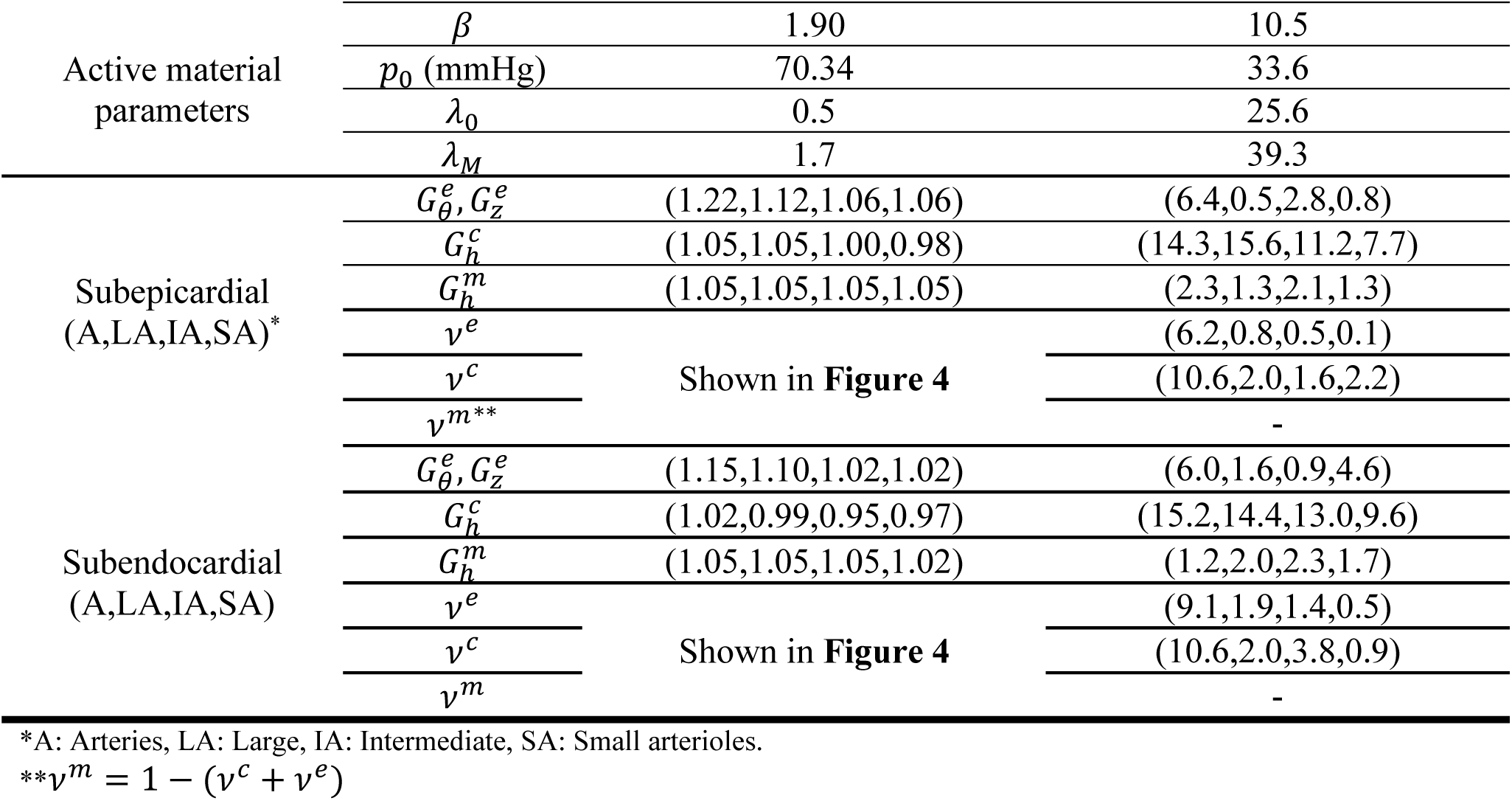
Constrained mixture model parameters, and their estimated value and sensitivity index. Experimental data shown in Fig. 5 were used for estimation of the parameters.

**Figure 4.**
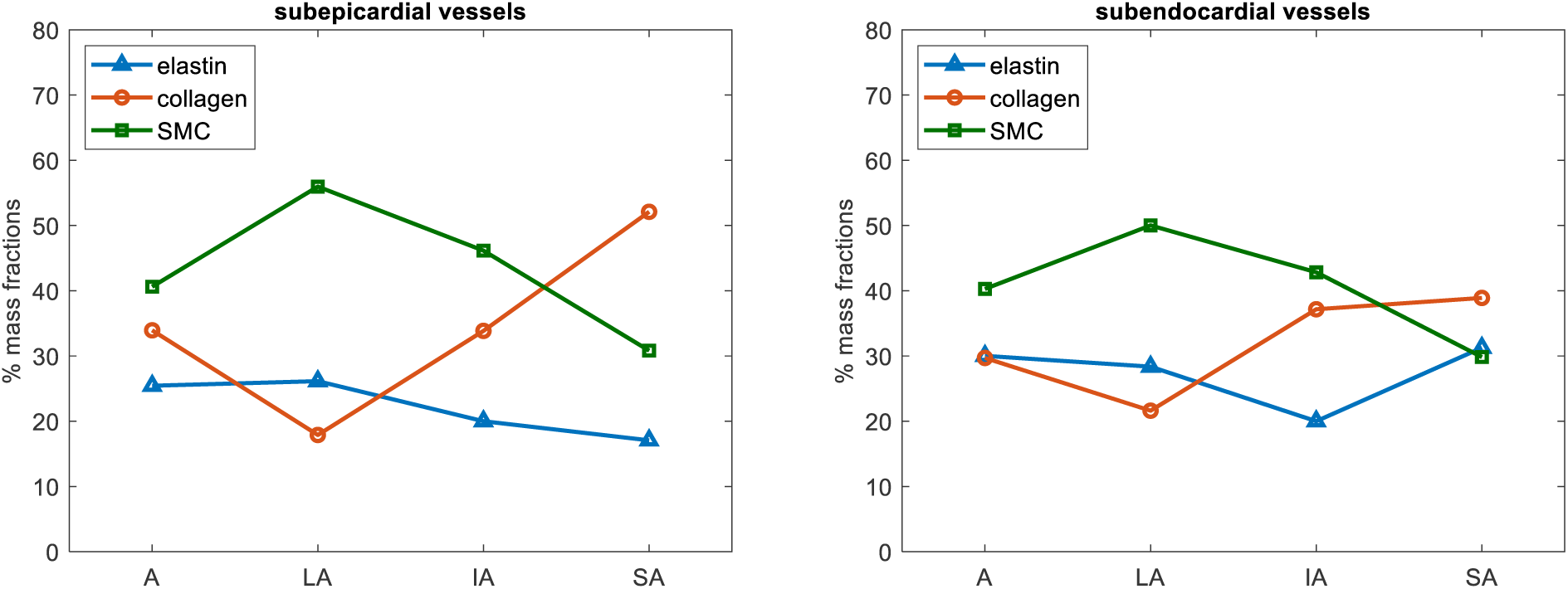
Estimated mass fractions (*v*^*e*^, *v*^*c*^, and *v*^*m*^) for the different vessel types in our model. The mass fraction of SMCs seems to be the highest in large arterioles in both subendocardial and subepicardial layers. (A: Small arteries, LA: Large, IA: Intermediate, SA: Small arterioles.)

**Figure 5.**
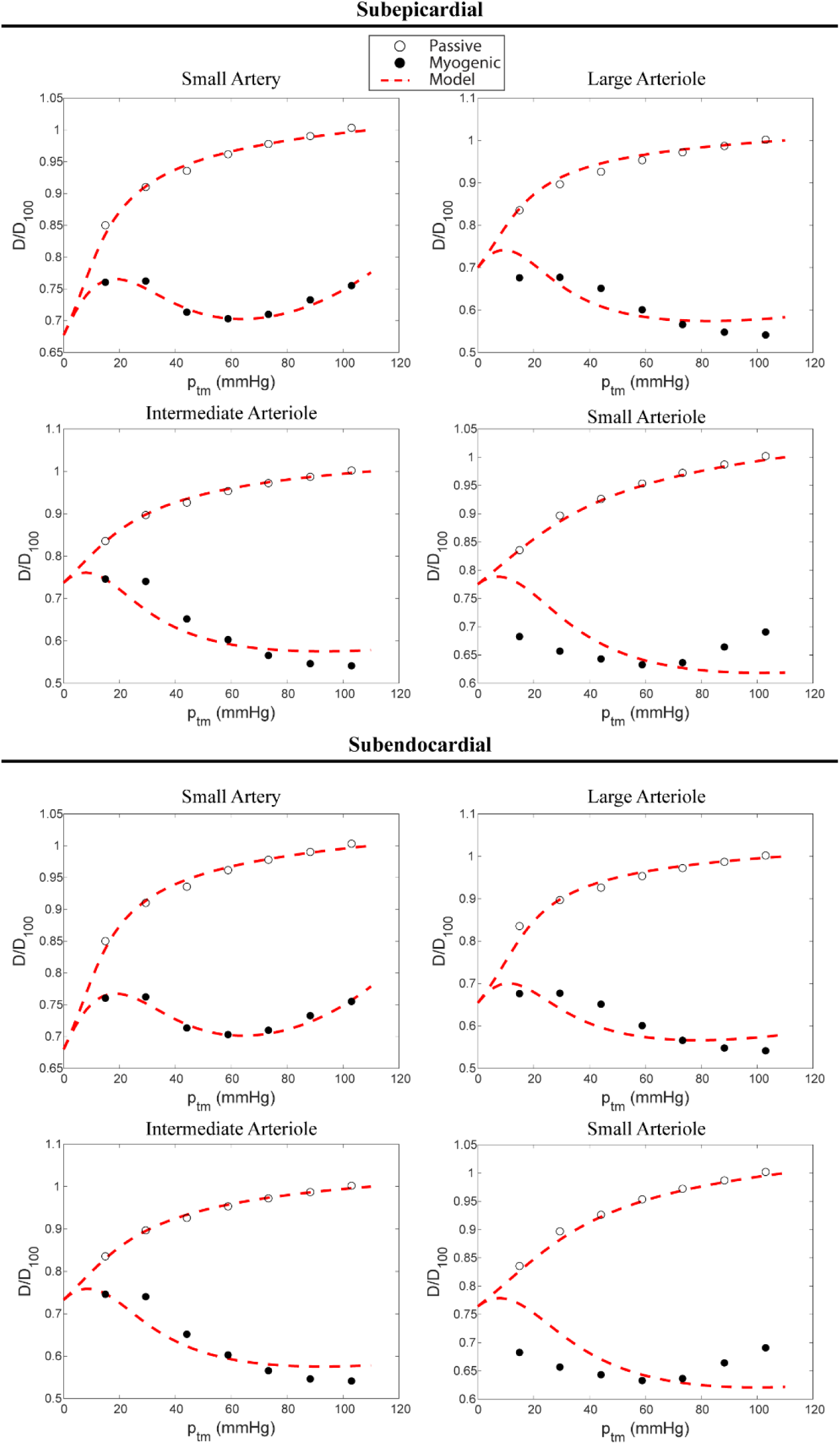
Data on passive and myogenic responses (55), and model fits for the representative vessels in the subepicardial (top) and subendocardial (bottom) trees. The parameters listed in Table 1 are estimated by minimizing the mean squared error between the predicted and measured diameters at the given values of transmural pressure estimate. *D*_100_: Passive diameter at 103 mmHg (100 cmH_2_O in the experiments.)

**Figure 6.**
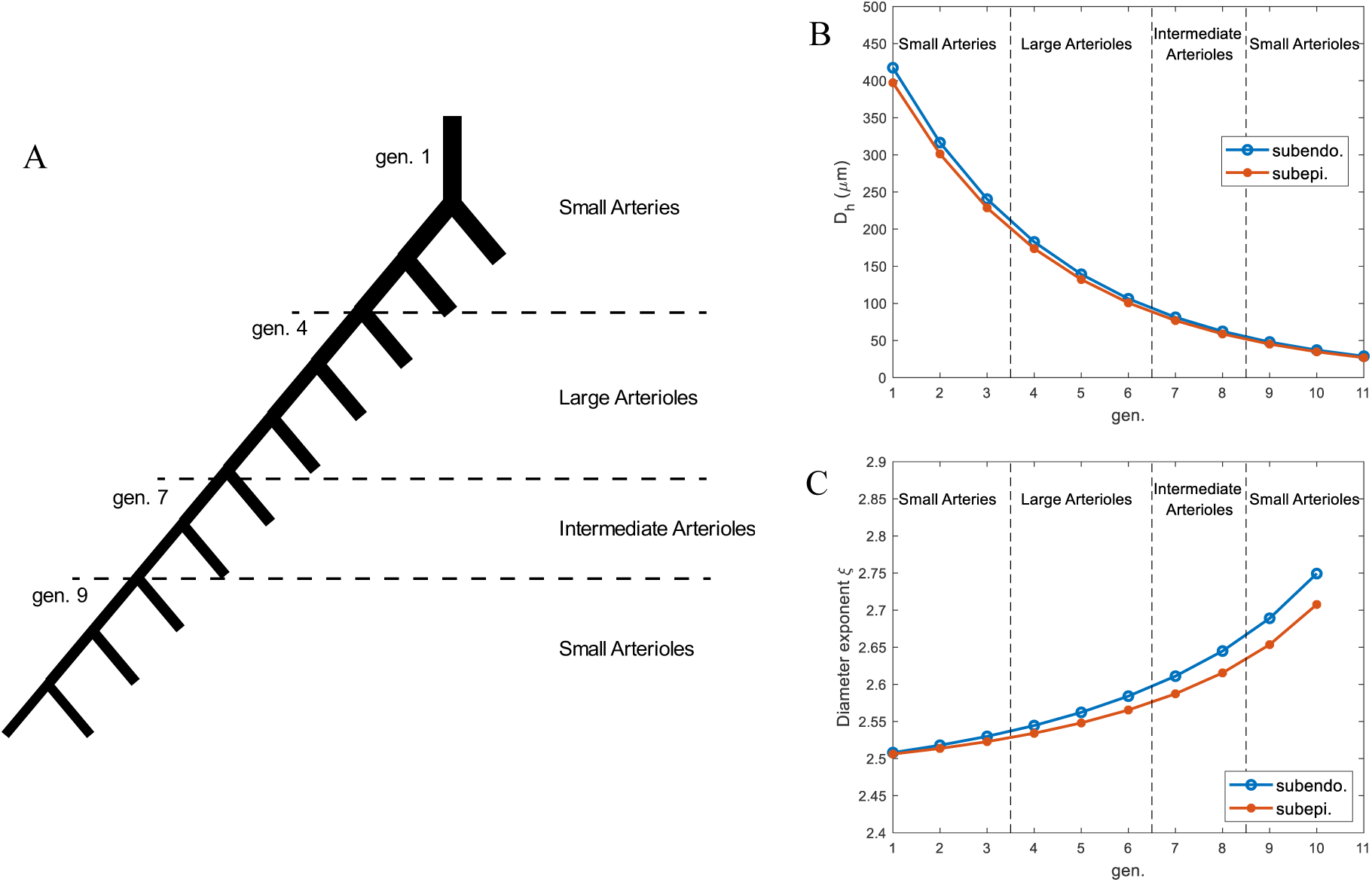
A: Schematic of a symmetric coronary tree, identifying the vessel types in each generation (number of generations and vessel classifications are the same in both subendocardial and subepicardial trees). B: Homeostatic vessel diameter D_h_ across generations for the subendocardial and subepicardial trees. C: Diameter exponent in daughter-to-parent diameter relation 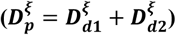 across generations for the subendocardial and subepicardial trees.

**Figure** 7 shows the structural and hemodynamics results of the homeostatic optimization for both subendocardial and subepicardial trees, and comparisons with available experimental data. **Figure** 7**-A** shows our results for thickness to diameter ratio for all generations. **Figure** 7**-B** shows pressure across the trees. It is apparent that most of the coronary vascular resistance is in arterioles < 100 μm. This range seems to be similar across different organs and species. Consistently, VanBavel and Spaan (86) showed that pressure in the coronary microvasculature drops from 90 to 30 mmHg in ∼10-μm diameter vessels. Wall shear stress increases almost 3-fold in arterioles, consistent with experimental studies (**Fig. 7-C**). **Figure 7-D** shows the homeostatic circumferential stress computed from Laplace’s law (*σ*_*h*_ = *p*_*tm*_*D*/(2*H*)), where *H* is the vessel wall thickness. Since the intramyocardial pressure is smaller towards the epicardium (resulting in higher transmural pressure), the subepicardial homeostatic circumferential stress is larger. Guo and Kassab (33) analyzed the circumferential stress in the swine coronary arterial tree and reported the circumferential stress in the range of 9.4-159 for arteries of 10-3,000 μm. Although our findings show the same trend, the Guo and Kassab experiments correspond to an entire coronary tree (not divided between subendocardial and subepicardial regions) and were conducted in an arrested heart with no intramyocardial pressure, therefore rendering larger transmural pressures and circumferential stresses.

**Figure 7.**
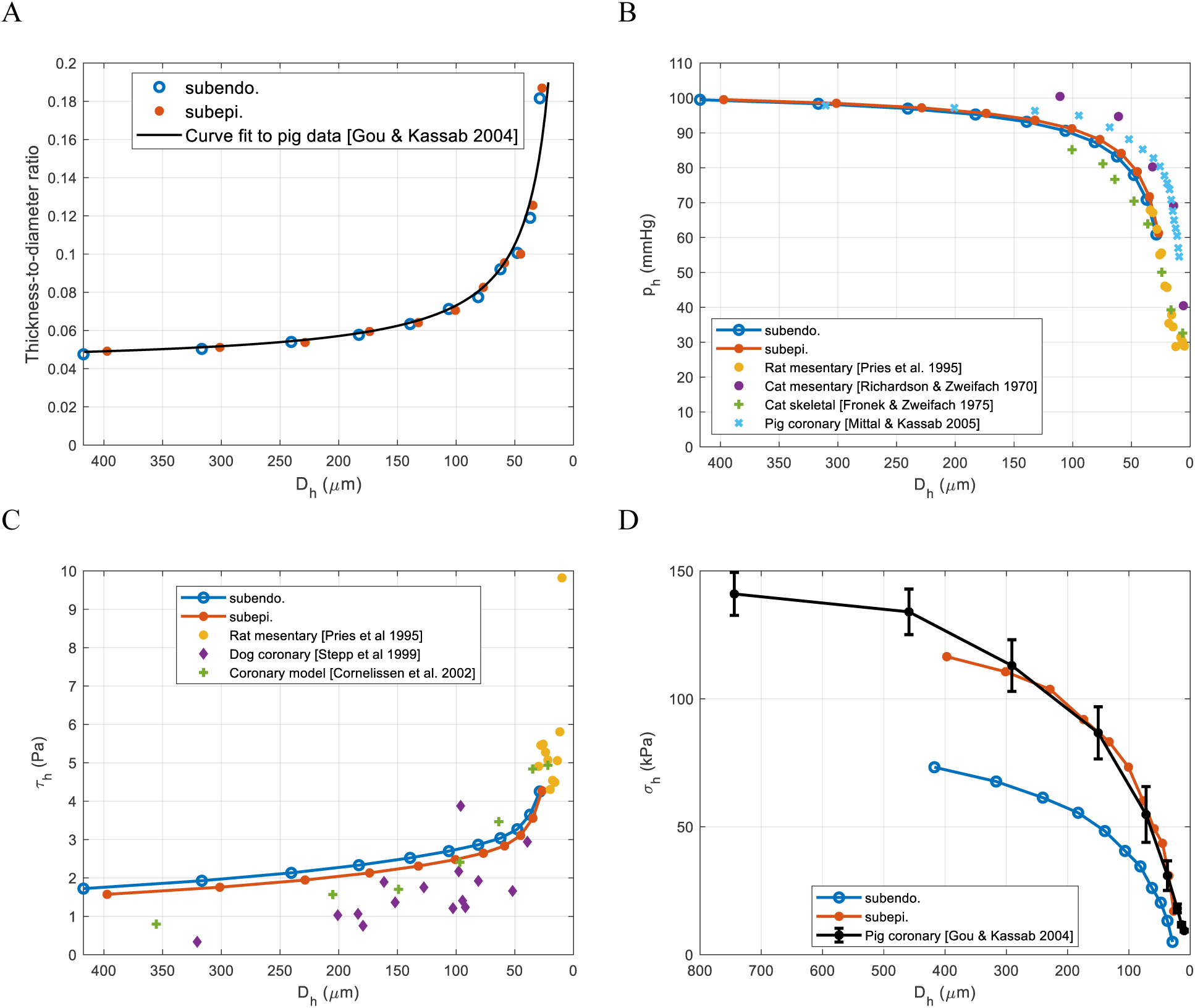
Homeostatic optimization results plotted against diameters: A: wall thickness-to-diameter ratio; The thickness to diameter ratio increases as the vessels become smaller. B: mid-artery pressure; Most of the coronary vascular resistance resides in arterioles < 100 μm. C: homeostatic value of wall shear stress (*τ*); A 3-fold in wall shear stress is predicted in the arterioles. D: homeostatic value of circumferential stress *σ*_*h*_; larger transmural pressures in the subepicardial layer compared to subendocardial layer result in larger homeostatic stresses in the blood vessels.

The homeostatic activation level (*A*_*h*_) and normalized active stress of the SMCs (*S*_*h*_/*S*_*max*_) across both coronary trees are shown in **Fig. 8**. The activation level decreases from the arteries to large arterioles (**Fig. 8-A**). A key outcome of the homeostatic optimization is that there is a trade-off between the SMC mass fraction and the activation level in the small arteries and large arterioles, where the pressure remains relatively constant. For instance, a higher SMCs content in the large arterioles (**Fig. 4**) results in a decrease in activation level compared to small arteries. However, for the intermediate and small arterioles, the transmural pressure decreases drastically across the vessel generations. The homeostatic activation shows different patterns for the subendocardial and subepicardial tress. Given that the SMC mass fractions are similar for both trees, we argue that the different activation level could be due to the substantially different transmural pressures due to the much larger intramyocardial pressures in the subendocardial region.

**Figure 8.**
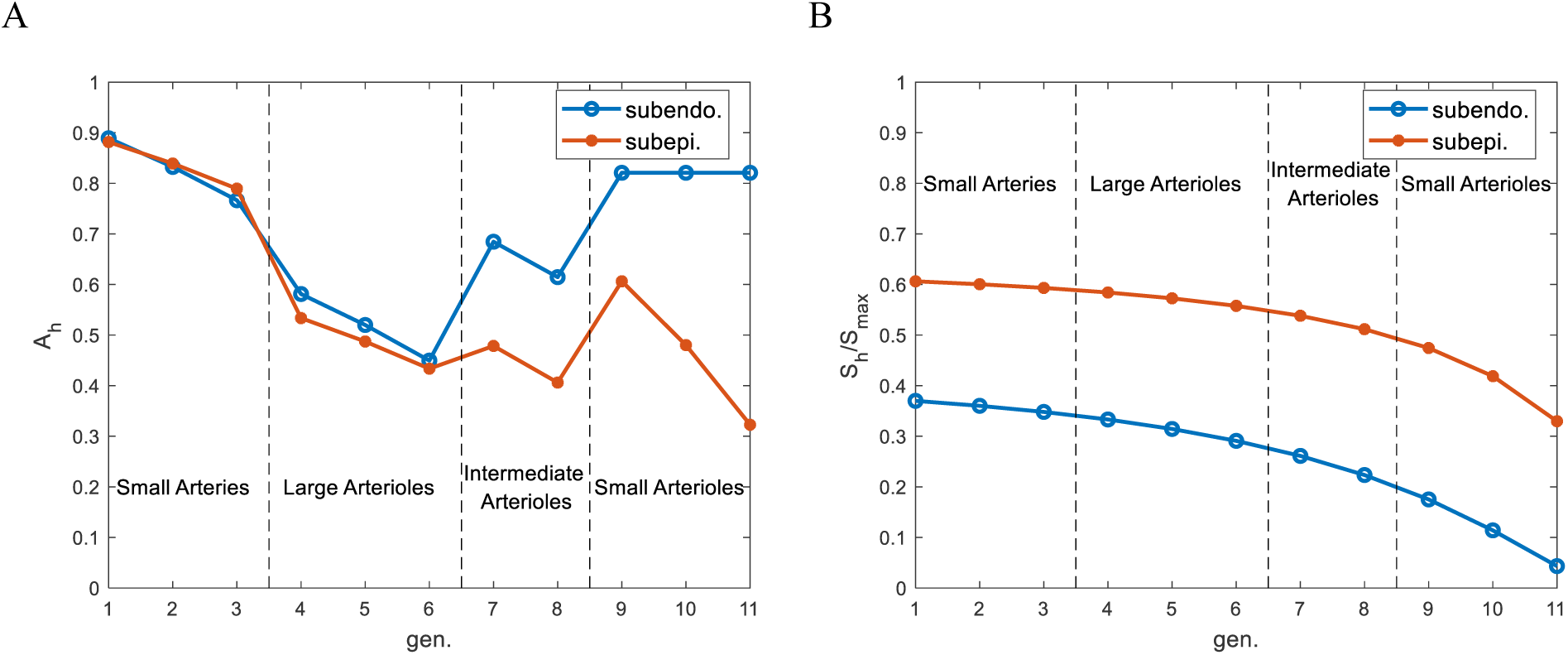
The homeostatic activation level (*A*_*h*_) and normalized active SMC stress (*S*_*h*_/*S*_*max*_) across both coronary trees.

Furthermore, a high level of baseline activation in the small subendocardial arterioles (∼0.8) is multiplied by a small (myogenic) pressure-dependent active stress *S*_*p*_ due to a small transmural pressure, and results in a small active SMC stress (*S*_*h*_/*S*_*max*_<0.2) (**Fig. 8-B**). Overall, the homeostatic active SMC stress *S*_*h*_ decreases due to large decreases in transmural pressure (**Fig. 8-B**).

### 3.3. Autoregulation model

Before presenting the results of the coronary autoregulation model, we must note that the outlet pressure prescribed in the homeostatic optimization was a constant pressure (55 mmHg) at the small arteriolar level. However, to extend the applicability of our coronary autoregulation model to situations of lower input coronary pressures (*p*_*in*_<55 mmHg), such as those distal to a severe occlusion in the large coronary arteries, we replaced the constant outflow pressure boundary condition with a constant resistance Rcoronary (with no vasoreactivity) boundary condition plus a 20 mmHg terminal coronary venular pressure (72). The values of these resistances were computed using the terminal arterioles pressure minus the venular pressure, divided by the terminal homeostatic flow in the coronary trees (viz., *R*_*coronary*_ = *ΔP*/*q*_*term*_).

#### 3.3.1. Calibration of stimuli, activation, and active stress

The scaling coefficients for the shear-dependent *a*_*τ*_ and metabolic *a*_*m*_ controls, were determined using swine pressure-flow autoregulation data (21) and a simplex method. As a constraint for the estimation of these parameters, we assumed that the SMC activation in small arteriolar level is less than 5% when coronary pressure falls under 40 mmHg (31). This physiologically motivated constraint helps to deal with high degree of underdetermination of the system. The estimated scaling coefficients are summarized in **Table 5**.

**Table 5.** Estimated scaling coefficients and their sensitivity index for the autoregulatory response. Experimental data shown in Fig. 9 were used for estimation of the parameters.

| | $a_\tau (X \%)$ | $a_m (X \%)$ |
| --- | --- | --- |
| Subepicardial | 1.03(0.3) | 10.34(1.0) |
| Subendocardial | 0.98(0.3) | 6.92(2.6) |

Figure 9 shows the results of the calibration of our coronary autoregulation model, together with the predicted pressure-flow relations in fully dilated coronary trees. The ratio of the total flow in the coronary arteries (*q*) over the baseline (homeostatic) flow (*qh*) is given as a function of the input coronary pressure *p*_*in*_. In the range of 70-140 mmHg inlet coronary pressure, over which coronary autoregulation is apparent, our model fits the experimental observations well (maximum error: 22% at 140 mmHg). Conversely, our model predictions are less accurate in the 20-60 mmHg inlet coronary pressure range (maximum errors of 43% at ∼50 mmHg inlet pressure). In low coronary pressures *p*_*in*_ < 60 mmHg, the predicted autoregulation curve is close to the fully dilated pressure-flow relation, indicating a full dilation of the trees.

**Figure 10** shows SMC activation level (*A*), pressure-dependent stress *S*_*p*_ (Eq. 3), and the SMC active tension *S* (Eq. 2) as a function of coronary pressure. Our results indicate that the activation levels in the intermediate and small arterioles increase monotonically with coronary pressure (**Figs. 10-A** and **B**). Conversely, the activation level remains relatively constant in the small arteries and large arterioles over the entire pressure range **Fig. 10**. The pressure-dependent stress *S*_*p*_ follows the same increasing trend with pressure in all the vessels in subepicardial tree (**Figs. 10-C**). However, this pressure-dependent stress is zero at low coronary pressure range in the subendocardium, since a large intramyocardial pressure in this layer (*p*_*im*_= 47 mmHg, cf. Section 2.3) causes a negative transmural pressure throughout the tree when the inlet coronary pressure is small (*p*_*in*_< 47 mmHg) (**Fig. 10-D**).

Active stress *S* is determined by the multiplication of the activation level and pressure-dependent stress *S*_*p*_ (Eq. 2). In low coronary pressures *p*_*in*_<50 mmHg, subendocardial vessels do not develop active stress *S*. Starting at pressure ∼50 mmHg the active SMC stress increases in the subendocardial tree with the coronary pressure (**Fig. 10-F**). In subepicardial arteries and large arterioles, the active SMC stress *S* is close to zero at *p*_*in*_= 20 mmHg but increases with coronary pressure. Conversely, it remains close to zero for small and intermediate arterioles (**Fig. 10-E**) for coronary pressures below ∼50 mmHg.

**Figure 9.**
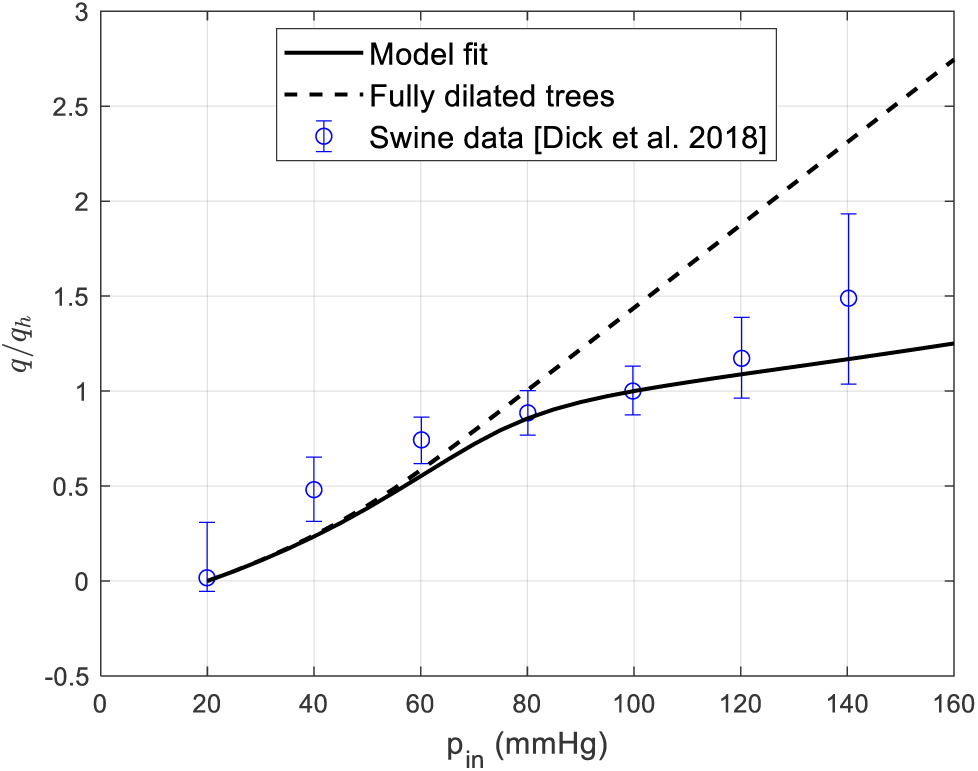
Pressure-flow autoregulation in combined trees compared to the collected experimental data in (21). The dashed line shows the pressure-flow relation when the entire tree is fully vasodilated. In the low coronary pressure range *p*_*in*_ < 60 mmHg, the autoregulatory and fully dilated responses are almost identical. *q*_*h*_ is the baseline flow rate with coronary pressure 100 mmHg.

**Figure 10.**
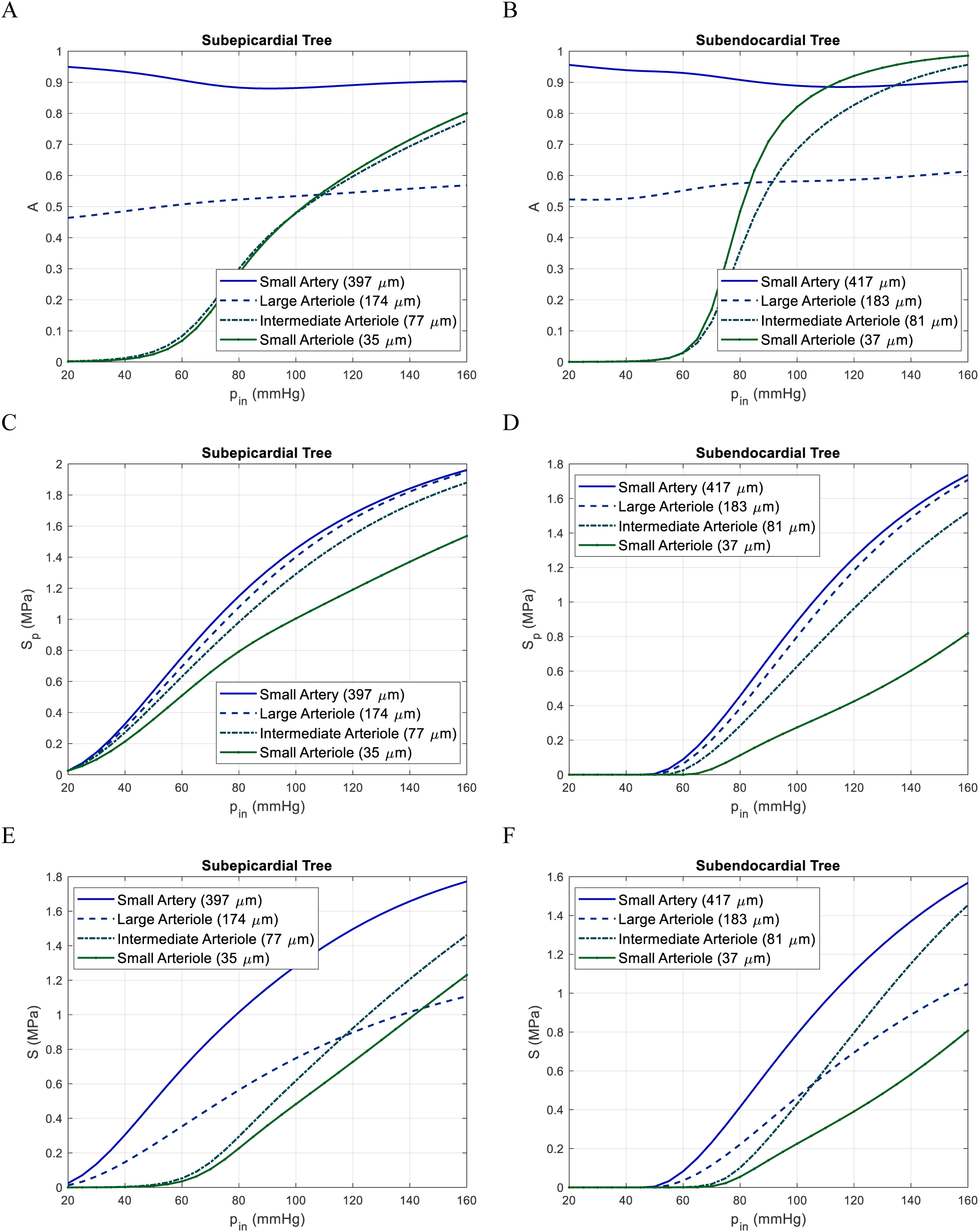
A & B: Activation as a function of the pressure at the inlet of the myocardial arterial. C & D: Pressure-dependent active stress *S*_*p*_. E & F: Active SMC stress *S*. Curves are given for all vessel types (small arteries, large, intermediate, and small arterioles) and both vessel tress: subepicardial (left column), and subendocardial (right column).

#### 3.3.2. Autoregulation responses for diameter, wall shear stress, and pressure

Figure 11 shows the variations of normalized diameters (*D*/*D*_*h*_), normalized wall shear stresses (*τ*/*τ*_*h*_), and normalized luminal pressure (*p*/*p*_*h*_), for four representative vessels in both the subepicardium and subendocardium trees as a function of the coronary pressure. Mediated by the autoregulatory mechanisms (myogenic, metabolic, and shear-dependent), blood vessel diameters D show a non-monotonic response to increase in coronary pressure in 20-160 mmHg range. Such behavior has also been observed in autoregulation of other circulatory systems such as the mesenteric circulation (15). Wall shear stress, however, presents distinct response patterns for the intermediate and small arterioles compared to the small arteries and large arterioles: For the intermediate and small arterioles, the wall shear stress increases monotonically with pressure. However, the wall shear stress remains near its homeostatic value in the *p*_*in*_ = 80-120 mmHg range for small arteries and large arterioles in the subepicardial layer (**Fig. 11-C)** and in the *p*_*in*_ = 100-140 mmHg range for small arteries and large arterioles in the subendocardial layer (**Fig. 11-D**). Pressure in both coronary trees monotonically increases with the coronary pressure (**Figs. 11-E** and **F**).

**Figure 12** shows the predictions of our model for the subendocardial to subepicardial flow ratio (ENDO/EPI) over the 20-150 mmHg pressure range. ENDO/EPI blood flow ratios have been reported between 1.09 to 1.49 across different species (22). Our results show that in low coronary pressures (*p*_*in*_ <57 mmHg), the ENDO/EPI ratio falls below one and they qualitatively match experimental data in the literature. For instance, Ball and Bache (11) reported a 50-60% decrease in the ENDO/EPI ratio in canine left ventricle as a result of mild to severe obstructions in large coronary arteries. Furthermore, Canty (14) indicated a redistribution of blood towards subepicardial tree (ENDO/EPI<1) for coronary pressures lower than ∼33 mmHg.

### 3.4. Model application: Response to drops in epicardial pressure

Figure 13 shows the predictions of diameter changes in our model for each coronary tree due to mild (*p*_*in*_ =59 mmHg) and severe (*p*_*in*_ =38 mmHg) reductions in coronary perfusion pressure, compared with experimental data (45) on epicardial vessels at the same pressures. Although the pressure reductions across stenoses are triggered by different levels of epicardial occlusion, our choices of inlet pressures are motivated by the experimental study by Kanatsuka et al. (45). For mildly reduced coronary pressures, the subepicardial arterioles (*D*_*h*_ < 190 μm) dilated, indicating a clear autoregulation response. Conversely, the subepicardial arteries remained near their baseline values, showing a mild vasoconstriction (**Fig. 13-A)**. Our results for the subendocardial vessels did not reveal substantial changes in diameter relative to their baseline values for the entire tree.

For a severe reduction in perfusion pressure, however, subepicardial vessels dilated in a similar fashion as in the mild reduction in pressure, while all subendocardial vessels constricted (**Fig. 13-B)**. This subendocardial tree constriction is likely due to the large intramyocardial pressures (*p*_*im*_= 47 mmHg), which in combination with the small luminal pressure (38 mmHg > *p* > 20 mmHg), results in a negative transmural pressure and therefore in vasoconstriction. The dilation of the subepicardial tree and constriction of the subendocardial tree explains the ENDO/EPI flow redistribution in low coronary pressures (**Fig. 13**).

**Figure 11.**
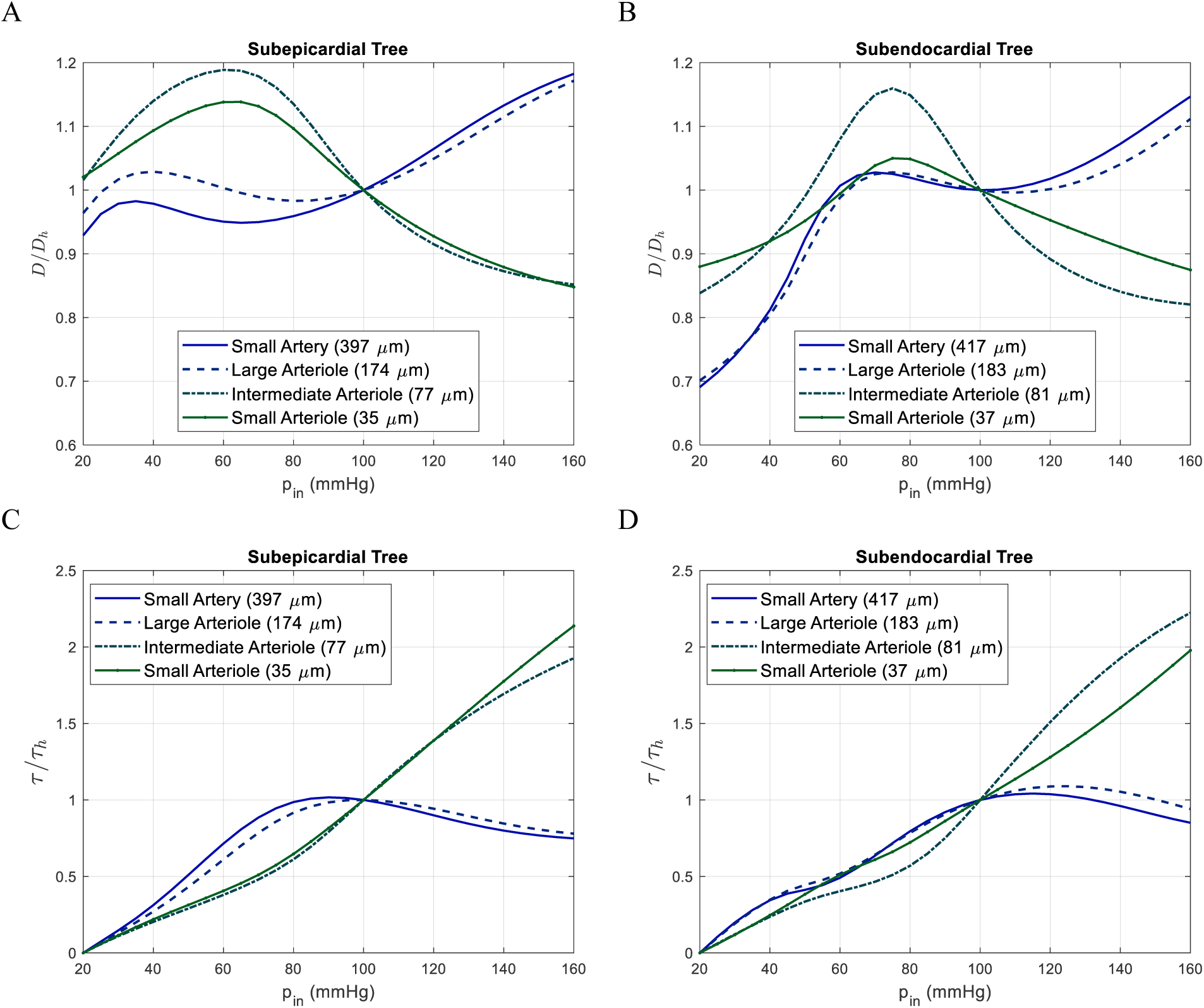
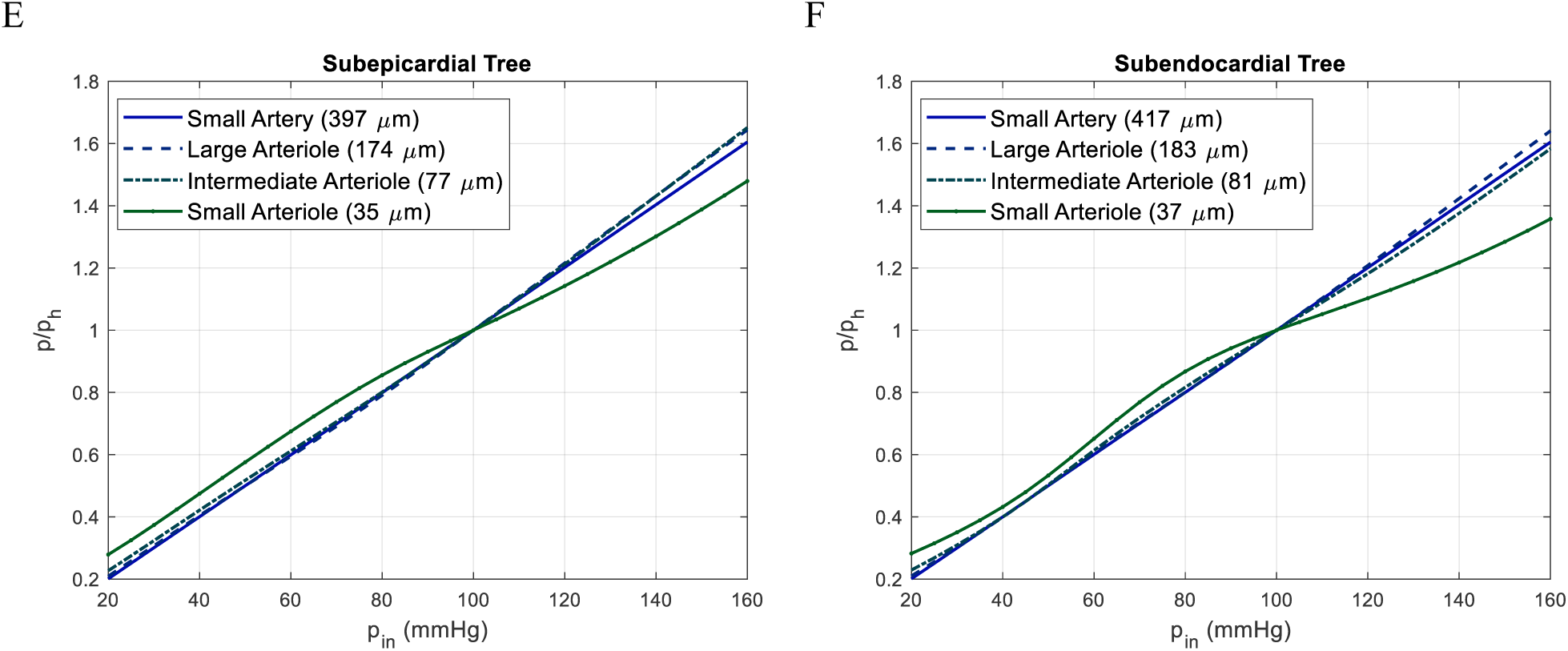
Normalized diameter *D*/*D*_*h*_, wall shear stress *τ*/*τ*_*h*_, and luminal pressure (*p*/*p*_*h*_) in four representative vessels as a function of the inlet coronary pressure. Vessel diameters *D* show a non-monotonic response to increases in coronary pressure. Some degree of regulation of wall shear stress in response to moderate changes in coronary pressure was predicted in small arteries and large arterioles in both layers. Pressure in vessels of both coronary trees increase as a function of the coronary pressure

**Figure 12.**
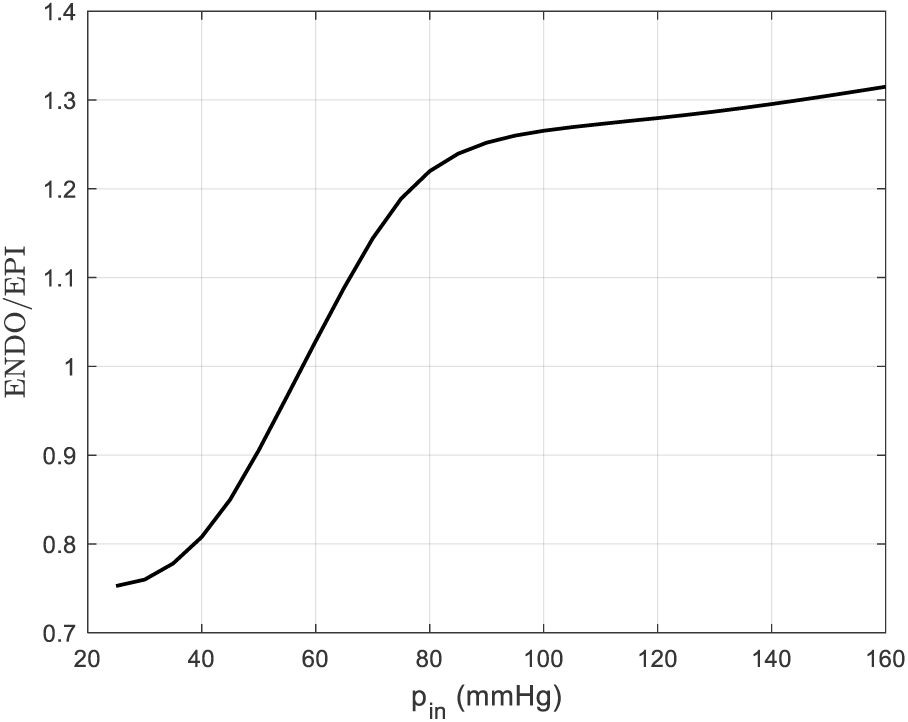
Transmural distribution of the blood flow during pressure-flow autoregulation. A regulation of the ENDO/EPI is predicted when the coronary pressure is 80-160 mmHg. In contrast, the ENDO/EPI drops significantly as the coronary pressure is reduced below 80 mmHg.

**Figure 13.**
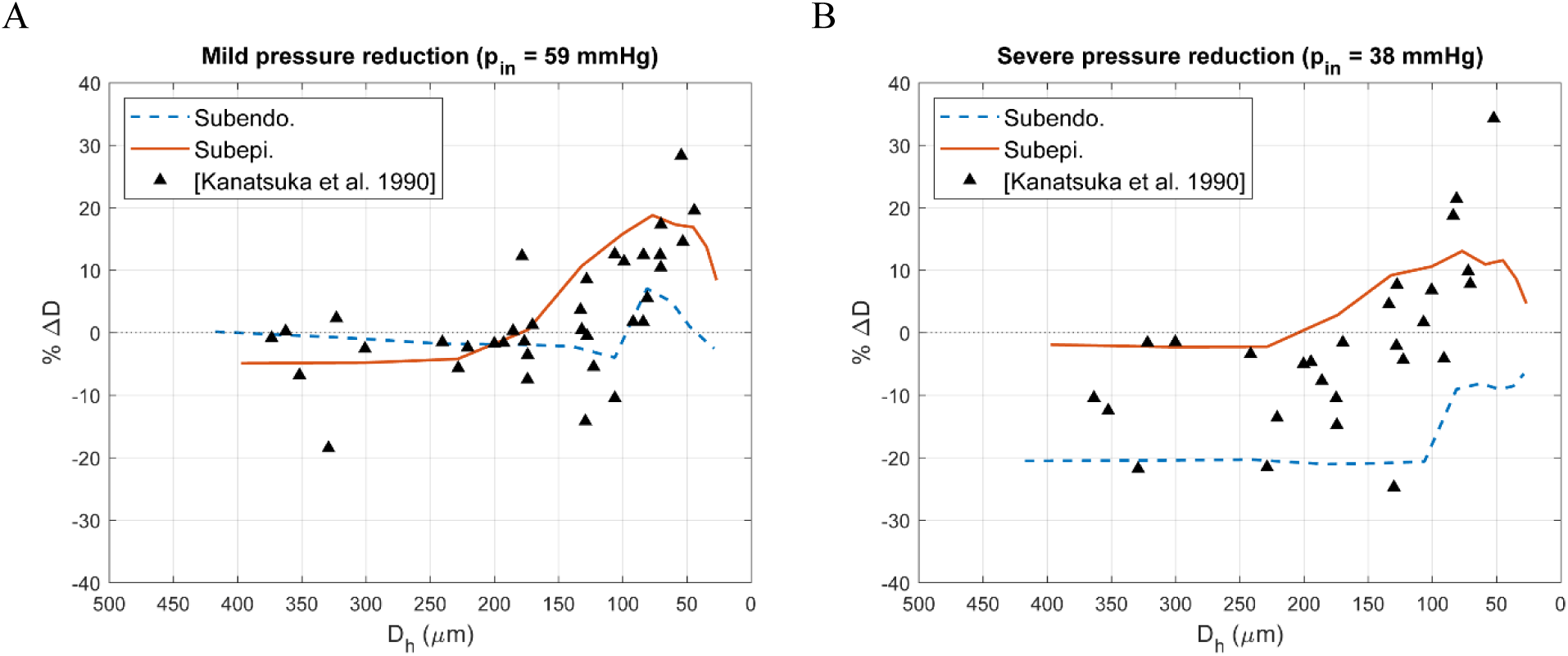
Effects of mild and severe pressure reduction on coronary arterial tree compared with experimental observations on canine epicardial vessels (45).

### 3.5. Model application: Effects of adenosine and L-NAME infusion

We further demonstrated our model by simulating responses of the coronary trees to two different pharmacological agents. **Figure 14** shows the model results together with experimental data (41) for diameter changes in response to intracoronary adenosine and L-NAME infusion.

**Figure 14.**
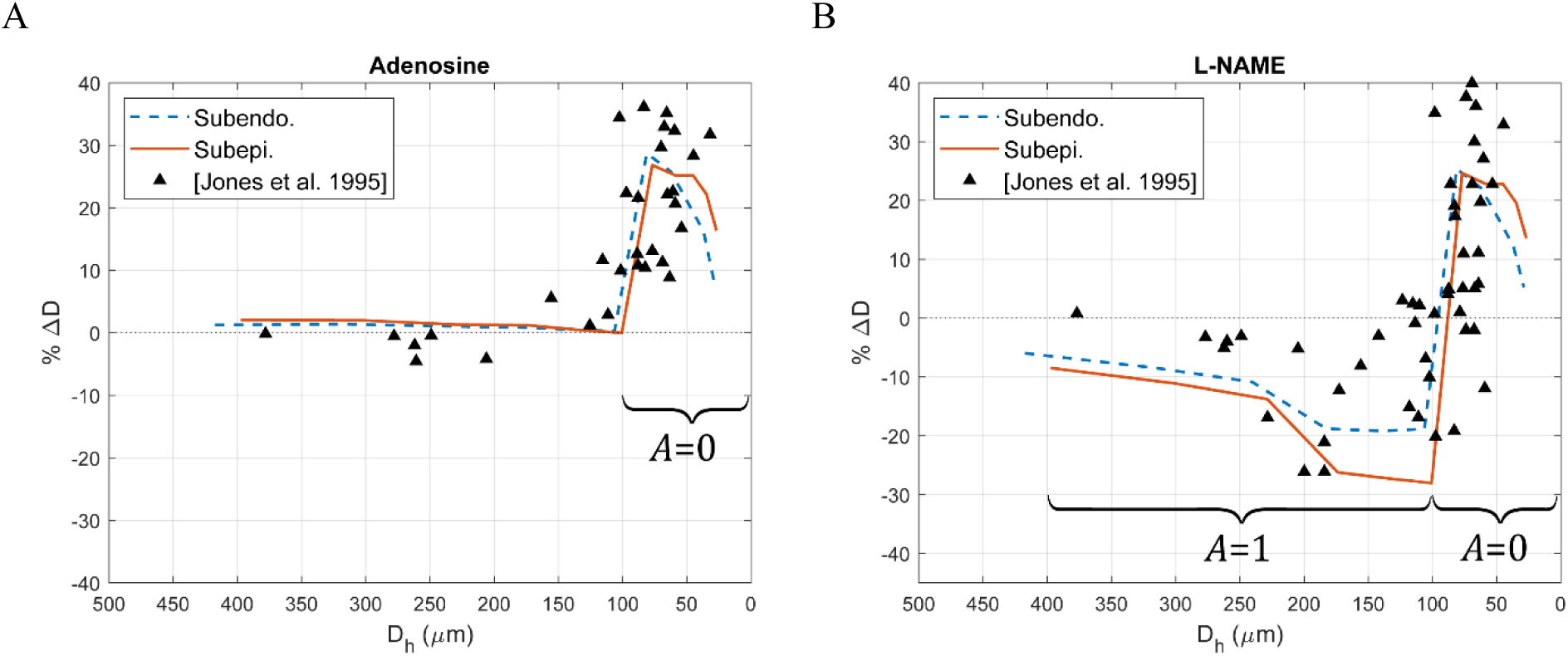
Effects of adenosine and L-NAME infusion on coronary trees compared with experimental canine data on epicardial microvascular response (41). Adenosine infusion was modeled by full dilation of small and intermediate arterioles *A* = 0. L-NAME infusion was modeled by full dilation of small and intermediate arterioles *A* = 0 and full myogenic constriction of large arterioles and small arteries *A* = 1.

Adenosine is a metabolic vasodilator which binds to purinergic receptors and causes SMC relaxation mostly in the intermediate and small coronary arterioles (55). In this work, adenosine administration was modeled by setting the activation in intermediate and small arterioles (*D*_*h*_ ≤100 μm) to zero (*A*=0). **Figure 14-A** shows that adenosine infusion leads to up to 25% increase in the diameter of these arterioles, while the diameters of small arteries and large arterioles remain close to their baseline.

L-NAME is an endothelial NO synthesis inhibitor (51). Based on the observations in the study by Jones et al. (41), L-NAME led to full myogenic constriction of small arteries and large arterioles (*D*_*h*_ >100 μm). However, they reported that although the experimental data showed both constrictive and dilatory responses in intermediate and small arterioles (*D*_*h*_ ≤100 μm), most of the arterioles were dilated. In addition, they observed that administering adenosine to L-NAME infused hearts did not further affect the dilated arterioles, indicating that these arterioles were likely fully dilated in response to L-NAME. Therefore, we set *A* = 1 for small arteries and large arterioles and *A* = 0 for intermediate and small arterioles. Our results revealed small arteries vasoconstriction in both trees and a higher degree of vasoconstriction in the large subepicardial arterioles compared to their subendocardial counterparts (**Fig. 14-B**). The latter can be due to a higher transmural pressure *p*_*tm*_ leading to larger myogenic pressure-dependent stress (Eq. 3). Diameter changes in intermediate arterioles (50<D≤100 μm) were similar in both coronary trees. Lastly, the small subendocardial arterioles (D≤50 μm) of were less dilated compared to the small subepicardial arterioles due to the larger intramyocardial pressure.

We must note that a direct comparison between model results and the experimental data on canine epicardial vessels is not feasible. This is due to the conceptual separation between subendocardial and subepicardial trees in our model, a separation not possible to establish in the experimental data. Furthermore, the experimental data in **Figs. 13** and **14** were acquired in dogs, whereas our model had been calibrated using porcine data. Nevertheless, our model managed to capture the overall responses to changes in perfusion pressure and administration of adenosine and L-NAME.

## 4. Discussion

The coronary circulation manifests a complex dynamic response to changes in perfusion pressure over multiple timescales. These timescales span from autoregulatory variations in active SMC tension, taking place within minutes, to slower changes in vessel microstructure (83) or vascular networks driven by angiogenesis or rarefaction over the course weeks to months (70).

Recent computational modeling endeavors in coronary physiology focused on capturing key features of short-term coronary autoregulation using lumped-parameter models (18, 66, 89). However, since these approaches lack structurally-motivated models for vascular tissue, they are not equipped for predicting long-term pathophysiologic responses of the coronary circulation. To address this need, this work aimed to produce a unified computational approach that could integrate vascular responses at the short time scales (e.g., autoregulation) as well as long time scales (e.g. vascular growth and remodeling) while including morphometric, hemodynamics and structural data. This design goal resulted in the adoption of a structurally-motivated modeling framework, widely used by the tissue growth and remodeling community, such as the constrained mixture theory. The constrained mixture enables integration of microstructural properties and cellular level functions within the vessel wall in a nonlinear continuum mechanics framework. The proposed autoregulation model was developed in three stages (**Fig. 1**). The main findings of each of the three stages of the model is discussed next.

### 1. Constrained mixture model parameter estimation

The parameters of the constrained mixture model were estimated using passive and myogenic pressure-diameter data (55). These parameters included passive and active material properties, constituent pre-stretches, and their mass fractions. Experimental observations have shown that in the physiological condition and in the presence of *in-vivo* levels of active SMC tension, the extracellular matrix components in the arteriolar wall are wavy and are likely under a state of compression (58). This observation was captured in our parameter estimation which rendered a collagen fiber pre-stretch in the small arterioles smaller than 1 (**Table 4**). Another determining factor in the *in-vivo* structure of coronary vessels is their myocardial location. The significant intramyocardial pressure on subendocardial vessels (47 mmHg) seems to compress collagen fibers even in large and intermediate arterioles (**Table 4**). Although the constituent pre-stretches are difficult to measure experimentally (37), they play a crucial role in determining the passive behavior of the vessel wall in its operating transmural pressure.

Furthermore, our model estimated a reduction in SMC mass fraction in the small arterioles relative to the large and intermediate arterioles (**Fig. 4**). This difference in estimated mass fractions seems to be due to the larger myogenic response in large and intermediate arterioles relative to small arterioles. A microscopic analysis of the mechanical structure of rabbit arterioles showed that the amount of SMCs gradually decreases from arterioles of 100 *μ*m to 30 *μ*m (75). Nevertheless, quantification of the mass fractions in both proximal and distal coronary arteries is essential for the mathematical modeling of functional (short-term) and structural (long-term) vascular adaptations in pathophysiological conditions (34).

Our sensitivity analysis showed that our model results are the least sensitive to the passive material parameters of elastin (*c*_1_) and SMCs (*c*_4_ and *c*_5_) (**Table 4**). Given that elastin is the main load bearing constituent at low pressures (88), the scarcity of pressure-diameter data in low pressure ranges (<20 mmHg) may contribute to this insensitivity. In addition, the passive response of SMCs may not be distinguishable from that of collagen fibers since they both contribute to the exponential stiffening of the passive pressure-diameter relations.

### 2. Definition of morphometry, structure, and hemodynamics in coronary trees

In the second stage of the model development, we used a homeostatic optimization to generate two symmetric coronary trees in subepicardial and subendocardial layers. This optimization (29) estimates the homeostatic morphometric (e.g, diameters), structural (e.g, thicknesses), and hemodynamics (e.g., wall shear stress) characteristics of a vascular tree by minimizing the metabolic and viscous energy dissipation under the constraint of mechanical equilibrium.

Two quantities were optimized in each vessel segment: 1) the homeostatic diameter and, 2) the total mass of wall constituents (elastin, collagen, and SMCs). **Figure 6-8** summarize the estimated homeostatic characteristics of the coronary trees. Our model predicted a consistent diameter exponent (2.5-2.75) compared to experimental data (**Fig. 6-C**). In particular, several studies on the morphometry of vascular trees have shown a varying diameter exponent *ξ* between 2 in larger vessels to near 3 in terminal arterioles (47, 60, 62, 79). Arts and Reneman (7) studied scaling laws in canine coronary vasculature and obtained an exponent of 2.55 for ∼400 μm diameter vessels. Suwa and colleagues (79) obtained an exponent of 2.7 by analyzing different vascular beds in several organs of the human body. Furthermore, Kassab and Fung (47) obtained an exponent of 2.73 for arterioles smaller than 50 μm in swine. In addition, our results for pressure, wall shear stress, and circumferential wall stress also showed reasonable agreement with experimental data as highlighted in **Fig. 6**. We must note that the inlet pressure for both subendocardial and subepicardial trees were assumed 100 mmHg based on the study by Mittal et al. (62) that showed the pressure does not significantly drop in vessels of 100-1000 μm. The predicted homeostatic activation level *A*_*h*_ seems to suggest that there exists a tradeoff between activation level and SMC mass fraction: a lower mass fraction of SMCs was associated with a higher activation for a given transmural pressure (**Fig. 8**).

### 3. Coronary autoregulation model

the proposed coronary autoregulation model was calibrated with experimental data on pressure-flow autoregulation. This is, to our knowledge, the first implementation of a constrained mixture theory to model coronary autoregulation. Our coronary autoregulation model is inspired by models previously proposed in the literature. For the myogenic control, we used the model proposed by (18), and for the shear-dependent and metabolic controls we used the activation model proposed by (15). This formulation was motivated by the ex-vivo experimental measurements in (55) where the transmural pressure controls the myogenic response. Alternatively, other modeling studies have assumed that the circumferential wall stress (*p*_*tm*_*D*/2*H*) sensed by the SMCs is the signal for the myogenic control mechanism (38, 87), based on earlier studies (40). This theoretical discrepancy is circumvented in our model since the dependence of myogenic response on the vascular wall thickness *H*, vessel diameter *D*, and the transmural pressure *p*_*tm*_ is implicitly integrated in the mechanical equilibrium equation. Furthermore, we designed our model such that we could separate the myogenic control from the other control mechanisms, to use the available myogenic pressure-diameter data. In addition, we formulated the shear-dependent and metabolic stimuli in terms of their deviation from a homeostatic value. This choice was motivated by previous applications of constrained mixture models in the study of long-term active SMC-mediated adaptations in cerebral vasospasms (10).

In this work, we showed that our model captures the essential pressure-flow coronary autoregulation over a pressure range of 70-140 mmHg (**Fig. 9**). However, our model showed larger deviations from experimental data for lower coronary pressures (20-60 mmHg). Over that pressure range, our model closely followed the fully passive curve. This behavior is explained by the patterns of active SMC stress *S* in **Fig. 10-E** and **F**, which revealed nearly no active tension in any vessels of the subendocardial tree and active tension present only in small arteries and large arterioles of the subepicardial tree. This behavior suggests that the larger vessels of the subepicardial region do not play a significant role in coronary autoregulation.

Our model produced different patterns of vessel dilation and constriction depending on vessel size and location within the myocardium (**Figs. 11-A** and **B**). As pressure reduces from baseline, *p*_*in*_=100 mmHg, towards *p*_*in*_=20 mmHg, small and intermediate subendocardial arterioles first dilate and then constrict (**Figs. 11-B**). The initial dilation of these vessels is driven by the autoregulatory response as seen by a decrease in activation *A* and active SMC stress *S* (**Figs. 10-B** and **F**). For coronary pressures <60 mmHg all the dilatory capacity in these vessels is exhausted (*A*∼0), and the reduction in transmural pressure leads to passive constriction of these vessels. Similar trends can be observed for intermediate and small arterioles in the subepicardial layer (**Fig. 11-A**). These vessels, however, always remain dilated compared to their baseline value as the coronary pressure is decreased to 20 mmHg.

In addition, an analysis of the size-dependent heterogeneity of the autoregulatory response in the moderate coronary pressure range (80-120 mmHg) reveals that the intermediate and small arterioles display more pronounced diameter changes in response to perturbations in coronary pressure, whereas larger vessels do not display similar variations (**Figs. 11-A** and **B**). Similar findings were reported in *in-vivo* epicardial coronary arterioles canine data (16), and feline and murine skeletal muscle arterioles (32, 61), where the proximal vessel diameters were almost irresponsive to pressure changes. The response of small arteries and large arterioles to changes in coronary pressure, however, is better understood in terms of the wall shear stress. Our results demonstrated that wall shear stress is regulated over a range of pressures (80-120 mmHg) at the level of arteries and large arterioles (**Figs. 10-C** and **D**), whereas wall shear stress is not regulated in smaller vessels. Similar observations were reported in *in-vivo* experimental observations on canine coronary vessels (78).

We demonstrated our autoregulation model in two different application examples: 1) responses to drops in epicardial pressure and 2) responses to adenosine and L-NAME infusion. The first application example showed reasonable agreement between simulation and experimental data. **Figure 12** shows that a severe reduction in pressure (e.g., severe stenosis) leads to a 20% diameter reduction of the proximal subendocardium vessels, rendering this layer susceptible to ischemia (4). This vulnerability was also evident in the redistribution of flow towards subepicardial layer in the pressure reductions <60 mmHg as shown in **Fig. 12**. Finally, our model was able to capture the experimental trends following adenosine and L-NAME infusion reported in (41) (**Fig. 14)**. These results demonstrate that our *in-silico* model could successfully integrate experimental data on morphometry and structure and then reproduce key patterns of coronary autoregulation.

The present study has several limitations. First, the experimental measurements delineating the pressure-diameter relationships of myocardial coronary vessels are scarce. We used four available sets of pressure-diameter data for a wide range of subepicardial and subendocardial vessels from 400 μm to 20 μm. More pressure-diameter data in the coronary arterioles will enhance the accuracy and predictive capabilities of the model. Although this limitation may not be influential in low coronary pressure ranges due to small activation, it may affect the total flow in high coronary pressures. In addition, the results of our constrained mixture model parameter estimation show a discrepancy with the myogenic pressure-diameter data in the small arterioles. In this work, coronary vascular networks were idealized to bifurcating trees with 11 generations, whereas a realistic reconstruction of the coronary network could impact the local hemodynamics and autoregulatory responses. Moreover, there is no cross-talk between the subendocardial and subepicardial trees in our model, whereas in reality the two layers of the coronary tree compete for flow in situations of low perfusion pressure. Our model was constructed based on steady state hemodynamics. However, hemodynamics in the coronary vessels are pulsatile. This pulsatility likely influences the homeostatic conditions of the wall constituents. Lastly, understanding the mechanisms involved in the coronary autoregulation, especially the role of metabolic control, has been a largely debated issue in coronary physiology (49). In this study we simulated coronary autoregulation using the three main mechanisms and their range of influence as defined in previous modeling studies. Our model, however, could be further refined with improvements in our understanding of the interplay between the different mechanisms of coronary autoregulation.

## 5. Conclusion

The study of physiology and pathophysiology in coronary circulation benefits from accurate mathematical models that can account for both the microstructure and physiological function of arterial networks. In this study, we presented a structurally-motivated coronary autoregulation model that uses a nonlinear continuum mechanics approach to account for the morphometry and vessel wall composition in two idealized coronary trees. Literature data were used to calibrate and test our model. With some modifications, this model can be applied to morphometry-based coronary trees instead of idealized symmetric trees. Finally, since our model is based on constrained mixture theory, it could be expanded to also study long-term growth and remodeling in the coronary circulation in response to hypertension, atherosclerosis, etc.

## ACKNOWLEDGEMENT

This study was supported by the NIH U01-HL135842 and R01-HL139813. We would like to also acknowledge Prof. Lik Chuan Lee and Dr. Vasilina Filonova for their valuable insights.

### Appendix

#### A. Mechanics of a single artery

A single segment of the arterial tree is considered as a thin-walled cylindrical tube composed of three main load-bearing constituents: elastin (*e*), collagen (*c*), and smooth muscle cells (SMCs; *m*). First, we only consider the passive response of constituents. Each constituent is assumed to separately contribute to the strain energy density:

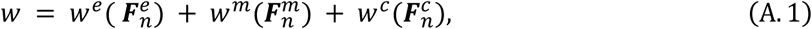

where 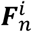 is the deformation gradient of each constituent *i* (*i* ∈ {*e, m, c*}) corresponding to its map from a stress-free configuration to the overall homeostatic configuration. We define this deformation gradient as 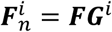, where ***F*** represents the deformation gradient of the mixture mapping from its reference configuration to the homeostatic configuration, and ***G***^*i*^ is a pre-stretch for each constituent mapping each constituent from its distinct stress-free configuration to the reference configuration (9). In particular, the pre-stretch mapping for elastin can be expressed as

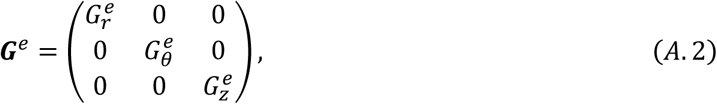

where 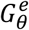 and 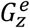 are pre-stretches associated with circumferential and axial directions, and 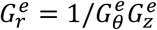.

Similarly, for collagen fibers and SMCs, ***M***^***i***^, *i* ∈ {*k, m*} is defined as the unit vector in the direction of the collagen fiber (*k*) or SMCs. The pre-stretch mappings for collagen and smooth muscle cells are given as

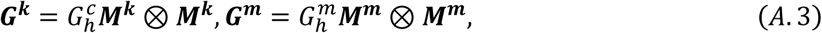

where the pre-stretches 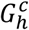 and 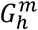 are also called homeostatic stretches, the stretches of the constituents when they are produced. We should note that in the previous applications of growth and remodeling, the pre-stretches were assumed to be constant for a single vessel. In our generalization of the framework to a vascular tree, we account for the variation of pre-stretches across the generations of vessels. Nevertheless, the pre-stretch implies that the homeostatic state in an individual vessel is associated with a constant homeostatic stress for the constituents of the vessel wall.

The orientation of collagen fibers and smooth muscles with respect to the axial direction in their reference configuration, defined by angle *γ*^*k*^, can be written as

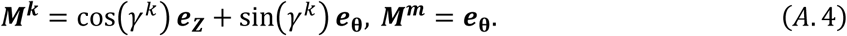

For modeling the extension and inflation of a thin wall model, ***F*** is considered as ***F*** = *diag*[*λ*_*r*_, *λ*_*θ*_, *λ*_*z*_]. The stretches of constituent *i, λ*^*i*^, *i* ∈ {*e, k, m*}, are expressed in terms of the pre-stretches using ***F***^*i*^ = ***FG***^***i***^

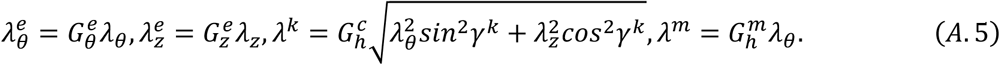

The incompressibility of the wall material is imposed by assuming an isochoric motion (i.e., det(***F***) = 1), and thus *λ*_*r*_ = 1/*λ*_*θ*_*λ*_*z*_. Using the membrane theory (36), the passive membrane Cauchy stress (force per deformed length) can be written as

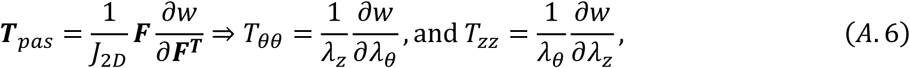

where *J*_2*D*_ = *λ*_*θ*_*λ*_*z*_. The total strain energy per unit area can be written as

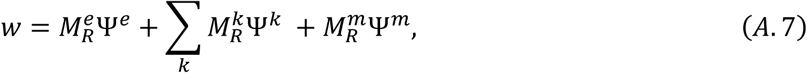

where 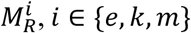 is the mass of each constituent per unit reference area. Alternatively, the total strain energy per unit area can be written as

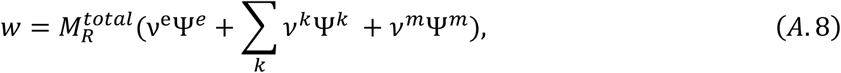

where ν^e^, *v*^*k*^, and *v*^*m*^ are mass fractions of elastin, collagen fiber families, and SMCs, respectively. In this work, four families of collagen fibers in circumferential, axial, and two diagonal directions were considered with mass fractions *v*^*k*^ =(0.1, 0.1, 0.4, 0.4)*v*^*c*^ where *v*^*c*^ is the total collagen mass fraction (90). The total mass per unit area 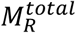 is the mass of load bearing constituents and can be computed via

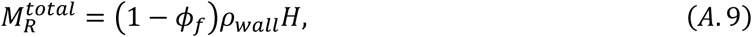

where *ρ*_*wall*_ is the density of the vascular wall, *ϕ*_*f*_ is the volume fraction of interstitial fluid, and *H* is the thickness under homeostatic conditions.

A neo-Hookean model is employed for the passive elastin response and a Holzapfel exponential model is used for collagen fiber families and passive behavior of circumferentially oriented SMCs

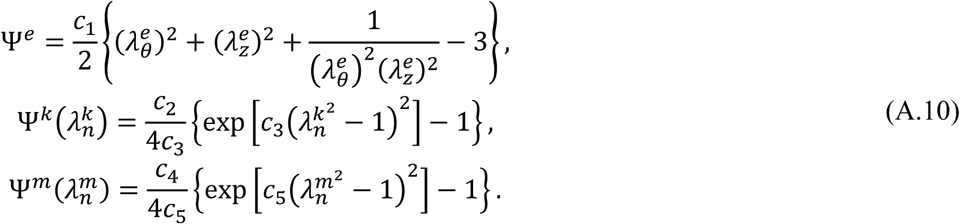

Where *c*_1_ is the elastin material parameter, *c*_2_ and *c*_3_ are collagen material parameters, and *c*_4_ and *c*_5_ are passive SMC material parameters. To include the active tension of vascular SMCs, we use a potential function as given by (10)

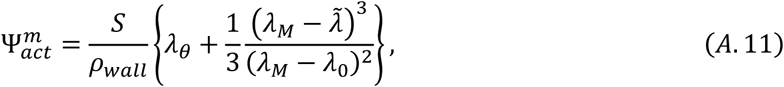

where *λ*_*M*_ and *λ*_0_ are stretches at which the active force generation is maximum and zero, respectively, and *ρ*_*wall*_ is the vascular wall density. In addition, 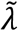 is an active stretch of the SMCs in the circumferential direction, which can evolve by SMC remodeling over slow timescales (hours to days). In the current study, we focus on the short timescale adaptations (minutes), and thus, 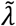 is assumed to be the total circumferential stretch in the vessel (*λ*_*θ*_). Active SMC stress *S* is in general a function of the myogenic, shear-dependent, and metabolic controls in coronary vessels. Again, using membrane theory, 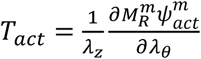 the total tension in the artery can be written as

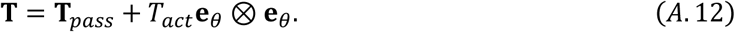

Finally, for a thin walled cylinder with transmural pressure *p*_*tm*_, the force equilibrium in the circumferential direction gives

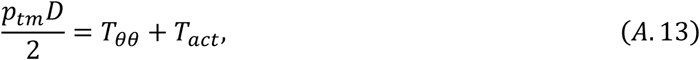

where *D* is the vessel diameter and *p*_*tm*_ is the transmural pressure of the artery.

We assume the reference configuration of the blood vessel in our continuum mechanics formulation is its homeostatic configuration. Therefore, by setting ***F*** = ***I*** (i.e., *λ*_*θ*_ = *λ*_*z*_ = 1 ) in equations above, we can write

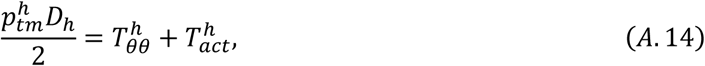

where 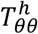 and 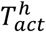 are homeostatic passive and active circumferential tensions, respectively, 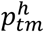 is the homeostatic transmural pressure and *D*_*h*_ is the homeostatic diameter of the vessel. The homeostatic passive tension 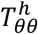 is thus determined by the material properties (*c*_1_-*c*_5_), constituent prestretches (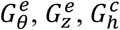, and 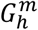) and their mass fractions *v*^*i*^. In addition, the homeostatic active tension 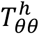 is determined by active SMC parameters, *λ*_*M*_, *λ*_0_ and *S*_*h*_. Accordingly, we can also compute the homeostatic stress in each constituent ***σ***^***i***^ as a function of its material parameters, its homeostatic stretch *G*^*i*^, and active SMC stress in case of SMCs.

#### B. Parameter estimation

The parameter estimations in sections 2.2 and 2.4 are performed by minimizing the following error function:

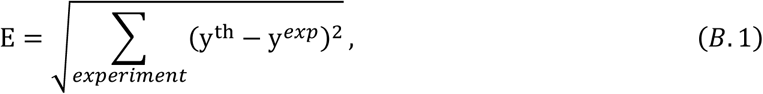

where *y*^*th*^and *y*^*exp*^ are theoretical and experimental values of the quantity *y*. In section 2.2, the quantity *y* is the diameter of vessels in passive and myogenic responses, and in section 2.4, the flow in the coronary arteries (*q*) over the flow when coronary pressure is 100 mmHg. Specific constraints for the optimization problem are stated in the text.

#### C. Homeostatic optimization

Originally introduced by Murray (64), and later extended by Taber (80) and Lindström et al. (56), the extended Murray’s law states that the vessel wall composition and geometry strive to minimize the energy consumption. The homeostatic optimization framework proposed in (29) leverages this idea to identify the homeostatic characteristics of an arterial tree. Briefly, the total energy cost per unit length *C* for an individual blood vessel can be written as

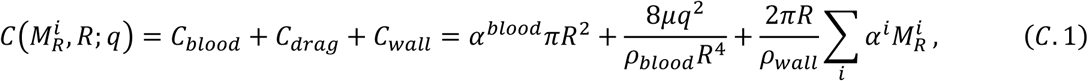

where *C*_*blood*_ is the metabolic cost of sustaining blood in the vessels, *C*_*drag*_ is the power needed to overcome viscous drag, and *C*_*wall*_ is the metabolic cost of maintaining the constituents of the wall. In addition, *α*^*blood*^ and *α*^*i*^ are the metabolic energy costs of blood and wall constituents (*i* ∈ {*e, k, m*}, see Appendix A) per unit volume, *R* (= *D*/2) is the radius of the vessel, *q* is the blood flow in the vessel, and *μ, ρ*_*blood*_, and *ρ*_*wall*_ are the blood viscosity, blood density, vessel wall density, respectively. We must note that the metabolic cost of SMCs can be written as the summation of the metabolic cost of their maintenance 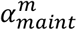 and the metabolic cost of the active SMC stress 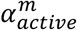.

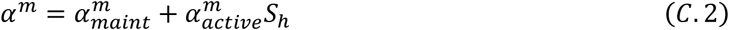

where *S*_*h*_ is the homeostatic active SMC stress. This metabolic cost function has to be minimized with Eq. A.14 as the constraint in each segment. With regards to Appendix A, we can rewrite the mass of constituents in terms of their mass fractions as 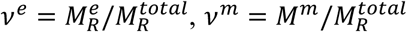, and 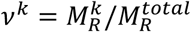 with *v*^*c*^ = ∑*v*^*k*^. Using the mass fractions and considering that the arteries are in homeostatic state, we can write the mechanical equilibrium in each vessel as

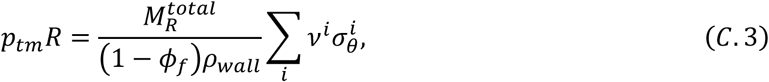

where *ϕ*_*f*_ is the volume fraction of interstitial fluid (70%) and 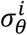 is the circumferentially acting part of the homeostatic stress tensor ***σ***^***i***^. Therefore, the energy cost of one vessel can be written in terms of diameter, with pressure and flow rate given from the hemodynamics as

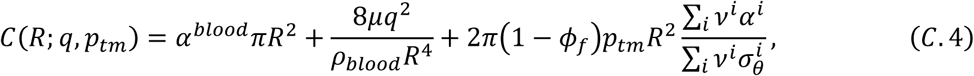

The total energy cost of all arteries can be added to compute the total cost for maintenance of an arterial tree

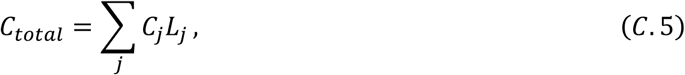

where *C*_*j*_ is the energy cost (Eq. B.3) and *L*_*j*_ is the length of segment *j*. The optimization requires the hemodynamics parameters such as *p*_*in*_, *p*_*out*_, and *q*_*total*_ as constraints. In this study, we fix the pressures at the inlet and outlets and incorporate the flow rate as a penalty term to the cost function.

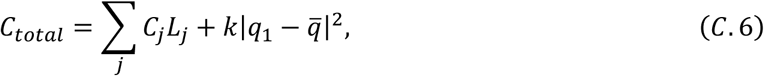

where *q*_1_ is the total flow in the tree, 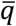 is the approximated flow in the tree from hemodynamic data, and *k* is a penalty coefficient. Once the radius of each vessel is determined by minimization of *C*_*total*_, the thickness of each vessel can be computed using equations B.2 and A.10. As explained in **Appendix A**, the homeostatic stress tensor ***σ***^***i***^ for each constituent is a function of its material parameters, its homeostatic pre-stretches, and the active stress *S* for SMCs. In this work, to find the homeostatic characteristics of the vessel wall in the tree, material parameters and pre-stretches are determined according to their diameter using the 4 types of vessels; small arteries, large, intermediate, and small arterioles. However, the SMC active stress *S* is tuned for each individual vessel in the tree by changing SMC activation *A* to fit the thickness to diameter ratio reported in Guo and Kassab (33). The parameters used for the homeostatic optimization are summarized in Table B1. thickness-to-diameter ratio with data in (33), as demonstrated in the results section.

**Table B1.** Parameters of the homeostatic optimization

| Parameter | Description | Value | Unit | Reference |
| --- | --- | --- | --- | --- |
| $\alpha^c$ and $\alpha_{maint}^m$ | Metabolic cost of collagen and SMCs per unit volume | 1070 | W/m<sup>3</sup> | (68) |
| $\alpha^e$ | Metabolic cost of elastin per unit volume | 0 (Only produced at early ages.) | W/m<sup>3</sup> | (56) |
| $\alpha^{blood}$ | Metabolic cost of blood per unit volume | 51.7 | W/m<sup>3</sup> | (57) |
| $\alpha_{active}^m$ | Metabolic cost of active SMC stress | 0.00872 | 1/s | (68) |
| $\rho_{wall}, \rho_{blood}$ | Vessel and blood density | 1060 | kg/m<sup>3</sup> | - |

#### D. Viscosity

Blood viscosity in the systemic vasculature depends on the size of the vessel and the hematocrit level (*H*_*D*_). Particularly, the variation of viscosity is more pronounced as the arteries and arterioles become smaller. In our study, we prescribe the viscosity using the following *in-vivo* viscosity law given in (73)

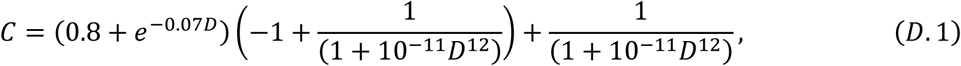

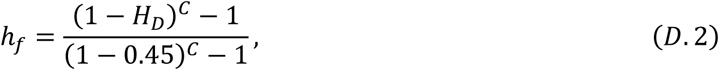

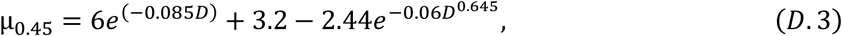

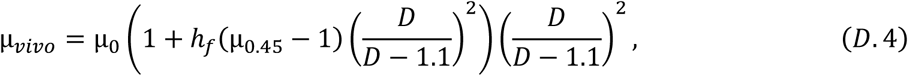

where *D* is the diameter in micron (2*R*), and μ_0_ is the viscosity of the blood plasma taken to be 0.001 Pa.s.

## Notes

### Competing Interest Statement

The authors have declared no competing interest.

